# Bayesian phylogenetics with incomplete lineage tracing data

**DOI:** 10.64898/2026.10.07.757283

**Authors:** Antoine Zwaans, Sophie Seidel, Kylie Chen, Tanja Stadler

## Abstract

Single-cell lineage tracing is rapidly emerging as a powerful method to track the dynamics of cell populations through time. Specifically, CRISPR-Cas9-based dynamic lineage tracing methods generate high-resolution data with the potential to investigate increasingly complex biological processes. However, single-cell readouts are inherently lossy, resulting in significant sparsity in lineage barcode alignments. Here, we present a mechanistic model of barcode loss implemented in the Bayesian framework BEAST 2, enabling phylogenetic inference from sparse datasets without the need for data filtering. Through simulations, we find that incorporating a mechanistic model of loss improves accuracy across all estimates in phylogenetic analyses by enabling previously unusable data to contribute to the inference. We further demonstrate the model’s utility across two datasets: one from a monoclonal cell culture and another from murine gastruloids. Consistent with our simulation findings, applying our model to these datasets recovers higher growth rates than workflows relying on standard data filtering, confirming that such filtering can lead to an underestimation of population expansion. Crucially, we demonstrate that our framework enables inference even from datasets whose sparsity previously prevented analysis, providing a robust foundation for extracting biological insights from lineage tracing experiments.

## 2 Introduction

Dynamic, genome editing-based lineage tracing technologies have emerged as powerful modalities to record relationships between individual cells in proliferating multicellular systems. Recent iterations of these technologies, leveraging targeted genome editing, have demonstrated the ability to record cellular histories at unprecedented temporal resolution and with high information density [1–5]. At the core of these approaches are artificial genomic barcodes that synthetically accumulate mutations as cell populations of interest grow, acting as recorders of cell lineages through time [6].

Sequencing a harvested cell population retrieves this encoded information, enabling the reconstruction of phylogenetic trees from barcode sequence alignments to recapitulate ancestral relationships among cells. When the branches of these trees are additionally scaled in time, they record the timings of cell divisions that lead to the final cell population under study. Such time-scaled phylogenies can be leveraged through phylodynamic inference to quantify crucial aspects of cellular proliferation and differentiation [7].

In classical phylogenetics, estimating time-scaled trees from sequence alignments relies on time-dependent mutation models operating under a molecular clock assumption [8]. Analogous to standard substitution models used for DNA sequence-based phylogenetic dating, mechanistic models describing genome editing can be integrated into maximum likelihood or Bayesian frameworks to infer time-scaled cell phylogenies from barcode sequence alignments.

Following this strategy, several specialized computational methods have recently been developed to accommodate specific features of single-cell and synthetic lineage tracing data. For instance, TiDeTree [9], GAPML [10], GABI [11], and LAML [12] are tailored to barcode-based technologies characterized by unordered mutation acquisition, such as GESTALT [13], IntMEMOIR [14], CARLIN [15], and BaseMEMOIR [5]. Conversely, SciPhy [16] was introduced as a software tool designed for technologies where mutations accumulate sequentially within barcodes, such as DNA Typewriter [2] and peCHYRON [4].

Beyond modeling genome editing mechanisms, a distinct hurdle in single-cell lineage tracing is accommodating the noise specific to single-cell sequencing read-outs. This noise consists primarily of sequencing errors and missing data. Missing barcode sequences can stem from sequencing dropout or, in methods relying on RNA readouts of transgenic constructs, from heritable epigenetic silencing that renders barcodes unobservable across entire groups of related cells. Together, these data loss mechanisms result in sparse sequence alignments. To accurately reflect these data-generating processes, mutation models can be extended to explicitly account for such single-cell error and loss patterns [17]. For somatic single-cell DNA sequencing data, this approach has been shown to improve accuracy in phylogenetic tree and divergence time estimates [18]. For synthetic lineage tracing data, the Bayesian framework TiDeTree [9] introduced a mechanistic model of barcode evolution for unordered edits [14, 19, 20], that accounted for a single source of data loss: heritable barcode silencing. While the maximum likelihood method LAML [12] extended this model to additionally account for a second source of loss–barcode dropout during sequencing–no existing Bayesian method accommodates both mechanisms. Furthermore, for sequential lineage tracing methods [2–4], no phylogenetic reconstruction method is capable of using lossy barcode alignments for inference, despite barcode loss being observed in all existing datasets published so far. For example, barcode loss ranges from less than 5% to up to 90% for DNA Typewriter datasets (see Fig. 1C).

**Figure 1.**
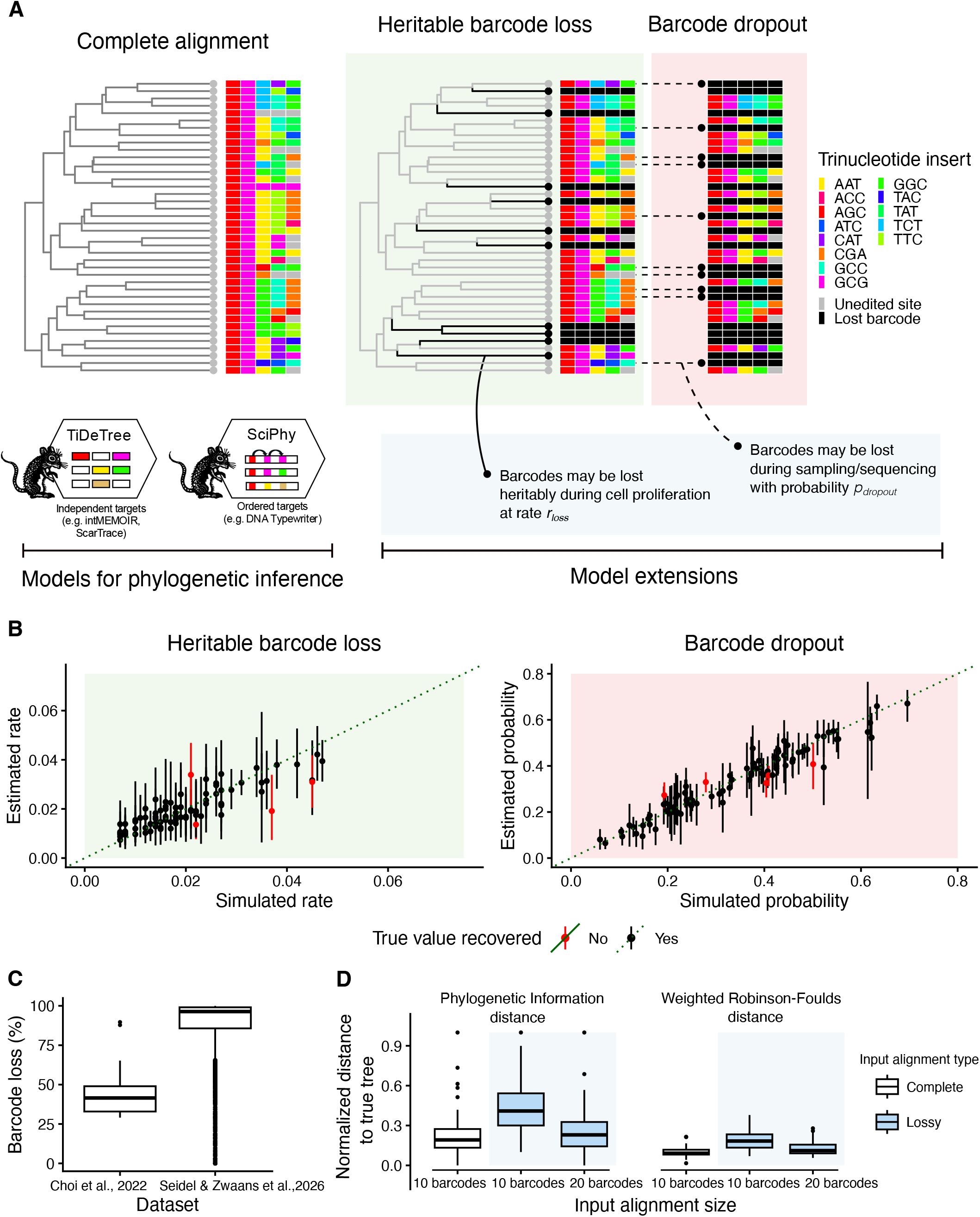
Mechanistic model of barcode loss and in-silico validation. **A** Comparison of idealized (complete, left) and realistic (lossy, right) lineage-tracing alignments simulated from a 40-cell population. We depict the two loss mechanisms incorporated in our model: ancestral heritable loss (e.g., transgene silencing, black branches) and stochastic sequencing dropout (dashed lines). **B** In-silico validation of the model extension for the SciPhy framework, heritable loss (left), and dropout (right). Dots represent posterior medians (black if the 95% HPD interval contains the true value; red otherwise) against a parity line representing the performance of a perfect estimator (green). **C** Empirical average barcode loss per alignment observed in published DNA Typewriter datasets. **D** Accuracy of tree reconstruction using simulated complete (left side of each subpanel) and lossy alignments (right side of each subpanel, in blue), of either 10 or 20 barcodes per cell. Performance is measured by distance to the true trees using Phylogenetic Information (left) and weighted Robinson-Foulds (right) metrics.

To address these gaps, we introduce a mechanistic model of barcode loss that enables the coherent use of sparse alignments within a unified Bayesian framework in BEAST 2. Building upon the existing Sci-Phy and TiDeTree packages [9, 16], we extend both frameworks to account for stochastic dropout and additionally introduce a heritable loss component to the SciPhy framework. Leveraging our implementation, we utilize simulations to investigate how phylogenetic estimates are affected by varying levels of alignment sparsity. Expectedly, compared to a complete alignment, a sparse alignment leads to less-resolved phylogenies. However, we show that standard filtering approaches, which discard samples and barcodes causing this sparsity, can introduce systematic biases into estimates of time-scaled trees, barcode editing dynamics, and population growth dynamics. Specifically, we show that these bias patterns are most pronounced for data obtained from populations characterized by high sampling proportions or significant heritable barcode loss. We demonstrate that our model extension—which operates unconstrained by filtering requirements and accommodates arbitrarily sparse datasets—corrects for these biases, thus enabling more robust phylogenetic inference. We further showcase the practical utility of our framework through the re-analysis of two published datasets using our extended SciPhy model: HEK293T monoclonal cell culture and murine gastruloids. Notably, in the HEK293T dataset, our mechanistic loss model yields significantly higher growth rate estimates than previous SciPhy analyses that ignored barcode loss, suggesting that the ignored sparsity may have previously led to an underestimation of expansion dynamics. Beyond correcting bias in high-quality datasets, the primary advantage of our framework lies in its ability to extract signal from sparse, traditionally unusable barcode alignments. By leveraging Shannon entropy to identify high-information barcodes, we successfully analyzed previously discarded murine gastruloid replicates, recovering phylodynamic signal that was previously considered inaccessible due to high sparsity. Ultimately, by introducing a coherent statistical accounting of data loss mechanisms to genome-editing models, our framework allows researchers to fully leverage sparse alignments without introducing filtering-induced biases, establishing a rigorous foundation for single-cell phylodynamic inference across lineage-tracing technologies.

## 3 Results

### In-silico validation

We introduce a unified Bayesian inference framework to account for sparsity in single-cell lineage tracing barcode alignments. Our framework explicitly decouples this sparsity, arising from barcode loss, into two distinct components: a heritable, irreversible loss rate during cell population growth (e.g., via transgene silencing) and a stochastic dropout probability during sequencing (see Fig. 1A). While a model accounting for heritable loss alone was introduced in TiDeTree [9] within BEAST 2, and a maximum-likelihood implementation incorporating both components was developed in LAML [12] for lineage barcodes characterized by unordered mutations, we unify the handling of sparse lineage tracing alignments within the BEAST 2 ecosystem for Bayesian inference [21]. Specifically, we extend TiDeTree with the dropout component and newly incorporate the complete two-component architecture into SciPhy [16] for barcodes characterized by sequential mutations. To integrate these developments, we re-parameterize the phylogenetic likelihoods of both TiDeTree and SciPhy (see Materials and Methods 5). This unified implementation formalizes the probability of observing a sparse alignment conditioned on the tree, a heritable loss rate, and a dropout probability across lineage tracing modalities analyzable in BEAST 2.

We confirm the correctness of our new implementations by conducting well-calibrated simulation studies [22] for both the TiDeTree and SciPhy extended frameworks. For each of them, we simulated 100 datasets under our loss model (see Tables 4 and 5 for parameter distributions) and subsequently estimated the phylogenies and parameters in BEAST 2 using priors that match the simulation setup. A Bayesian simulation study is considered well-calibrated if the true parameter value is contained within the *x*% credible interval in approximately *x*% of the replicates, a proportion known as coverage.

Our validation study suggests that both model extensions are well-calibrated. For the SciPhy extension, the 95% credible intervals for the heritable barcode loss rate and the dropout probability contained their true values in 95% and 94% of the replicates, respectively (see Fig. 1B). Similarly, the TiDeTree validation yielded well-calibrated estimates across parameters, achieving a coverage of 96% for the dropout probability and 94% for the heritable loss rate (see Appendix Fig. 6). All coverage values fall within the expected binomial range for a 95% coverage target [23], indicating that the new likelihood extensions are correctly implemented.

We observe a high correlation between simulated (true) parameters and their posterior estimates for both the heritable barcode loss rate and the dropout probability. For the SciPhy implementation, we achieved Pearson’s *R* values of 0.85 (95% confidence interval (CI) of [0.78, 0.90]) and 0.96 (95% CI [0.94, 0.97]), respectively. Similarly, the TiDeTree implementation demonstrated strong recovery with Pearson’s *R* values of 0.79 (95% CI [0.70, 0.86]) and 0.87 (95% CI [0.82, 0.92]). These results indicate that our framework effectively extracts signal for both mechanisms of barcode loss across a broad range of parameter values (see Fig. 1B and Appendix Fig. 6).

Having confirmed the validity of our implementation, we next evaluate the impact of data loss on phylogenetic reconstruction accuracy, focusing on the SciPhy model as a representative case, given that both models performed comparably in our validation study. Using the same sets of true trees from our validation, we compared the reconstruction accuracy obtained from complete (theoretically ideal) datasets against that obtained using (more realistic) lossy alignments as input. Predictably, for an equal number of barcodes, trees reconstructed from lossy data are consistently less accurate than those obtained from complete alignments. This performance gap is apparent in both topology and branch lengths, as quantified by Phylogenetic Information (PI) [24] and weighted Robinson-Foulds (wRF) distances [25] between the inferred trees (Conditional Clade Distribution (CCD) summary trees [26]) and the true simulated trees (Fig. 1D). Specifically, under our simulation setup, which leads to an average loss of *∼*50% of barcodes (driven equally by heritable loss and stochastic dropout; see Table 4), we find that achieving a reconstruction accuracy with lossy data (Fig. 1D, blue) comparable to that of complete data (white) requires at least double the number of barcodes; namely, 20 lossy barcode alignments to reach a similar accuracy as 10 fully recovered barcodes with identical editing dynamics.

### Extended simulation study

We next benchmark our unified framework against standard filtering practices to evaluate the impact of explicitly modeling data loss on phylogenetic inference. In the absence of formal, model-based frameworks to handle lossy data, single-cell phylogenetic pipelines predominantly rely on adhoc filtering, both removing cells with missing barcode sequences and discarding entire barcodes with incomplete alignments, to artificially construct “complete” datasets for inference [2–4, 16] (see Fig. 2A, left). We investigate how this heuristic filtering approach impacts phylogenetic accuracy compared to our newly implemented mechanistic loss model (see Fig. 2A, right). To ensure a fair comparison, we fixed the number of cells across both methods. Notably, while our mechanistic model offers the advantage of utilizing arbitrarily large, unfiltered datasets containing more cells (bounded only by computational limits), we intentionally downsampled its input to match the exact cell count retained after the filtering pipeline; this ensures that any observed accuracy gains are strictly driven by our handling of data loss rather than a larger sample size (see Materials and Methods for filtering details).

**Figure 2:**
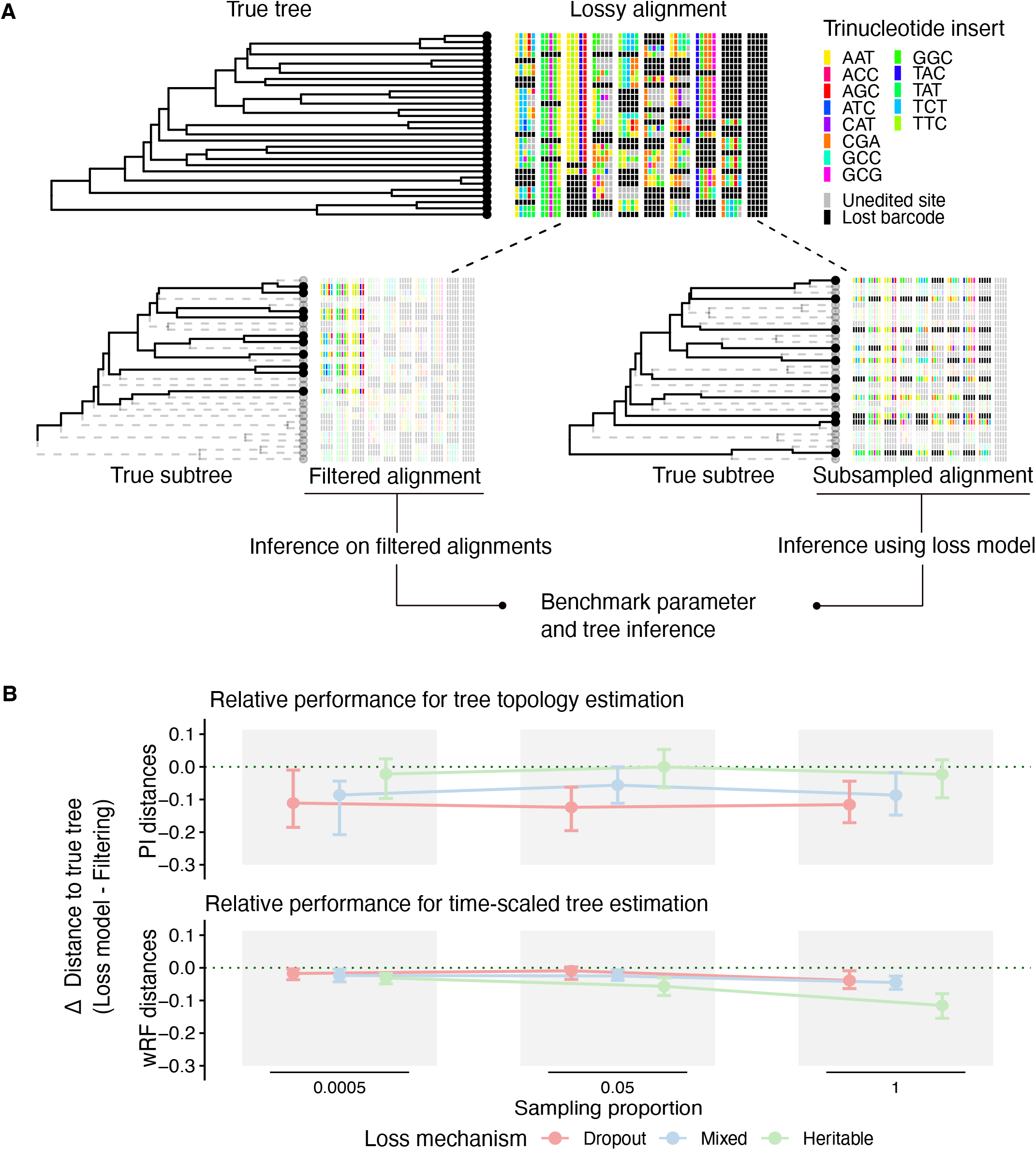
Simulation-based comparison of data filtering versus mechanistic loss modeling for tree estimation. **A** Schematic of data processing workflows. The traditional heuristic pipeline (left, “Filtered alignment”) discards cells with missing barcode sequences or removes entire barcodes with incomplete alignments to yield artificially complete datasets. In contrast, our framework (right, “Subsampled alignment”) retains these lossy entries, integrating them directly into phylogenetic inference via a unified mechanistic loss model. For a controlled benchmark in our simulation study, the unfiltered alignment is randomly subsampled to match the exact cell count of the “Filtered alignment,” generating the “Sub-sampled alignment” analyzed by our loss model. In empirical applications, however, our framework enables analyzing complete single-cell datasets without subsampling, bounded only by computational and runtime resources. **B** Comparative accuracy of tree reconstruction across loss scenarios (colors) and tree shapes (controlled by the sampling proportion). The y-axis shows the difference in distance to the true tree (*d*_Loss_ *™ d*_Filtering_), where negative values indicate higher accuracy for the mechanistic loss model. Topological accuracy is measured by Phylogenetic Information distance (PI; top), and time-scaled branch-length accuracy is evaluated by weighted Robinson-Foulds distance (wRF; bottom). Points represent median differences across 50 simulated datasets per condition; error bars denote the interquartile range.

Using both the filtering pipeline and our mechanistic model, we evaluate the recovery of the phylogenetic tree, editing rates, and population growth rates across three barcode loss configurations (dropout only, heritable loss only, and a combined model featuring both mechanisms, hereafter referred to as ‘mixed’) and three sampling regimes. These regimes were selected to test our hypothesis that differing branching patterns result in distinct signatures of data loss; accordingly, we modulated tree shapes by varying the sampling proportion *ρ* from low (*ρ* = 0.0005) to intermediate (*ρ* = 0.05) and complete (*ρ* = 1.0) sampling. This follows the logic that, for a fixed population size, lower sampling proportions push the observed branching events further back in time (closer to the root), whereas higher sampling captures more recent, terminal divisions (see Appendix Fig. 10). In total, we simulated 50 trees and alignments for each of these nine distinct combinations, allowing us to evaluate performance across a broad spectrum of both tree topologies and loss mechanisms.

As shown in Figure 2B, our mechanistic loss model consistently outperforms the traditional filtering approach across all loss scenarios and sampling proportions for the estimation of both tree topology and time-scaled phylogenies. However, the magnitude of this improvement follows opposite patterns depending on the accuracy metric used. In terms of topological accuracy, measured by the PI distance (Fig. 2B, upper panels), the mechanistic model provides the most substantial improvement in the barcode dropout scenario, with a median reduction in PI distance of approximately −0.12 across all sampling regimes. By comparison, the improvement is more modest for heritable loss, where differences are minimal (e.g., −0.0003 at intermediate sampling). The mixed loss scenario consistently falls between these two extremes. This hierarchy (dropout, mixed, heritable) is stable regardless of the sampling proportion. Conversely, the accuracy of estimated time-scaled phylogenies measured by the wRF distance (Fig. 2B, lower panels) reveals the opposite trend: the mechanistic model provides the greatest relative benefit under heritable barcode loss. In this case, we additionally observe a clear interaction with sampling intensity; as the sampling proportion increases, the advantage of the mechanistic model becomes more pronounced. This is most evident under heritable loss, where the median difference improves from −0.031 at low sampling (*ρ* = 0.0005) to −0.115 under complete sampling (*ρ* = 1.0).

In phylodynamic analyses, time-dependent parameter inference relies on extracting signal from branch length distributions in jointly inferred phylogenetic trees. To evaluate how potential inaccuracies in estimated branch lengths (reflected in the previous wRF distance results) impact the simultaneous quantification of the birth, death, net growth, and molecular clock rates, we assessed the accuracy of their posterior estimates via the absolute bias (Fig. 3A) and their precision via the relative HPD width (Fig. 3B). While estimates generated with our mechanistic loss model remained generally unbiased across all conditions, the filtering approach exhibited significant sensitivity to heritable loss. This sensitivity manifested as a strong positive bias in both birth and death rates that grew with the sampling intensity. At a sampling proportion of *ρ* = 1, for instance, the median bias reached 4.98 for birth and 5.08 for death rates, translating into a significant negative bias for the net growth rate (−0.20). Precision followed a similar pattern of degradation; while the mechanistic model maintained stable relative HPD widths across all loss regimes and sampling proportions (0.69–0.84 for birth rate, 2.73–3.30 for death rate), filtering led to severe loss of precision under heritable loss, reaching median relative widths of 7.82 for birth and 32.14 for death rates at *ρ* = 1. These results demonstrate that the mechanistic model not only protects against parameter bias but also preserves the precision of phylodynamic estimates. Similar patterns were observed for the clock rate, where median bias remained near zero for the loss model but reached −0.019 for the filtering approach under heritable loss at *ρ* = 1 (see Appendix Fig. 7).

**Figure 3.**
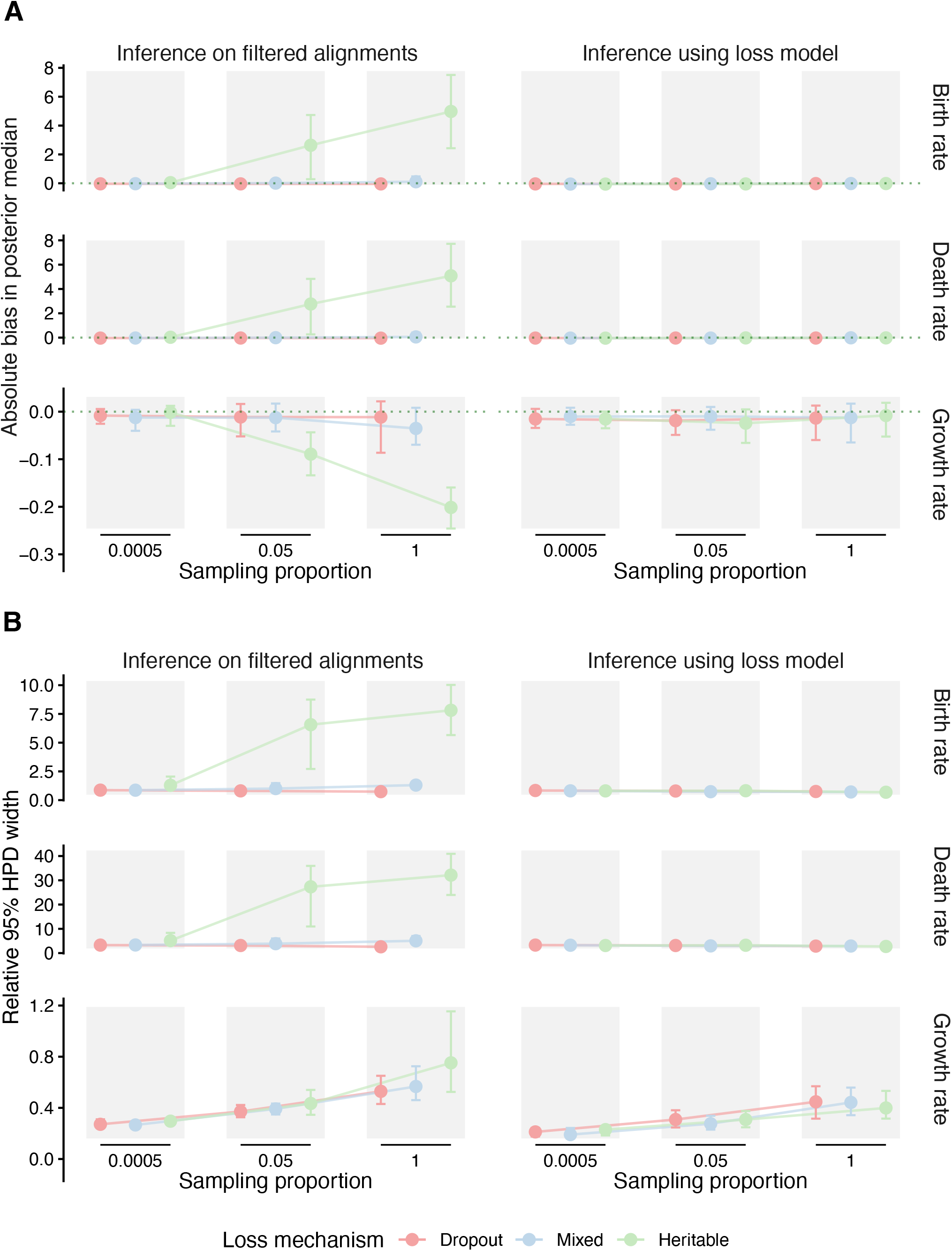
Simulation-based comparison of phylodynamic parameter estimation under data filtering and mechanistic loss modeling. **A** Absolute bias and **B** Mean relative HPD width for birth, death and net growth (birth − death). The estimation is compared varying loss scenarios (colors) and tree shapes (modulated by sampling proportion). Points represent the mean performance across replicates, with error bars denoting the 95% interval of the estimates. In A, the horizontal dotted line indicates an ideal bias of zero.

To investigate why heritable loss specifically compromises parameter estimates under the filtering approach, we examined the relationship between estimation bias and different features of barcode loss. By simulation design, the total percentage of barcode loss remained stable across all mechanisms and sampling proportions (mean *∼*50%; Appendix Fig. 8A), which allowed us to test whether bias is a function of loss magnitude or loss structure. Surprisingly, neither approach showed significant correlations between bias and the overall percentage of missing barcodes (*p >* 0.05; Appendix Fig. 8B), suggesting that the volume of loss is not the primary determinant of this distortion. We therefore sought to characterize the structural nature of the loss by quantifying the size of affected monophyletic clades. Heritable loss impacts significantly larger clades than stochastic dropout, a disparity amplified at higher sampling intensities (Appendix Fig. 9A). This occurs because higher sampling results in shorter pendant branches, increasing the probability that an ancestral loss event is inherited by multiple sampled cells (see Appendix Fig. 10). We identified the size of lost clades as the primary driver of systematic bias in the filtering approach, with strong correlations observed for both birth and death rates (e.g., *R ≈*0.79, *p <* 10^−30^ at *ρ* = 1; Appendix Fig. 9B). These results identify the removal of barcodes in large clades—rather than independent, unclustered loss—as the underlying cause of the observed bias. In contrast, the mechanistic loss model exhibited no significant correlations between clade size and bias (*p >* 0.05), demonstrating its robustness to this structured data sparsity.

### Application to previously analyzed lineage tracing datasets

To test if our simulation-derived insights hold true in an empirical setting, we applied our new model extension in SciPhy to a dataset of 25-day monoclonal expansion of HEK293T cells, tracked using the DNA Typewriter lineage tracing construct. In contrast to the analyses of [2] and [16], which were forced to exclude four of the 17 available barcodes due to incomplete alignments, our loss model allows us to utilize all 17 barcodes. To compare the resulting inference outputs, we generated a random sample of 1,000 cells and repeated the phylodynamic analysis from [16], simultaneously inferring the phylogeny, the editing dynamics of the ‘DNA tapes’ (barcodes), and the population growth rate, while additionally estimating our model’s loss parameters. In this analysis, we hypothesized that while the random sample leads to overall sparser data (see Appendix Fig. 14, mean loss of 46.2% over 17 barcodes), utilizing all available barcodes may avoid some of the biases identified in our simulation study. Figure 4 summarizes the tree and parameter estimates obtained using the random alignment sample in combination with our loss model. Interestingly, our estimates for the growth rate are consistently higher than those inferred when filtering incomplete alignments, a pattern that holds true regardless of whether the sampling proportion is fixed or estimated (see Fig. 4C and Appendix Fig. 13). These results are in close agreement with our simulation study (see previous section) which demonstrated that the filtering approach tends to underestimate net growth rates in the presence of heritable loss. The higher growth rate estimated under the loss model (posterior median 0.595 *d*^−1^, versus median of 0.540 *d*^−1^ with filtering), corresponds to a doubling time of approximately 28.0 hours (95% HPD: 25.6–30.4 hours), which is in precise agreement with the reported 24–30 hour doubling times for this cell line ([27]). Notably, this correction occurred despite a low inferred sampling proportion (posterior median *ρ ≈* 0.00009, Appendix Fig. 12D), a regime where our simulation study suggested the filtering approach should remain relatively robust. This suggests that the nature of barcode loss in the HEK293T expansion may be more severe than the stochastic mechanisms assumed in our simulation study.

**Figure 4.**
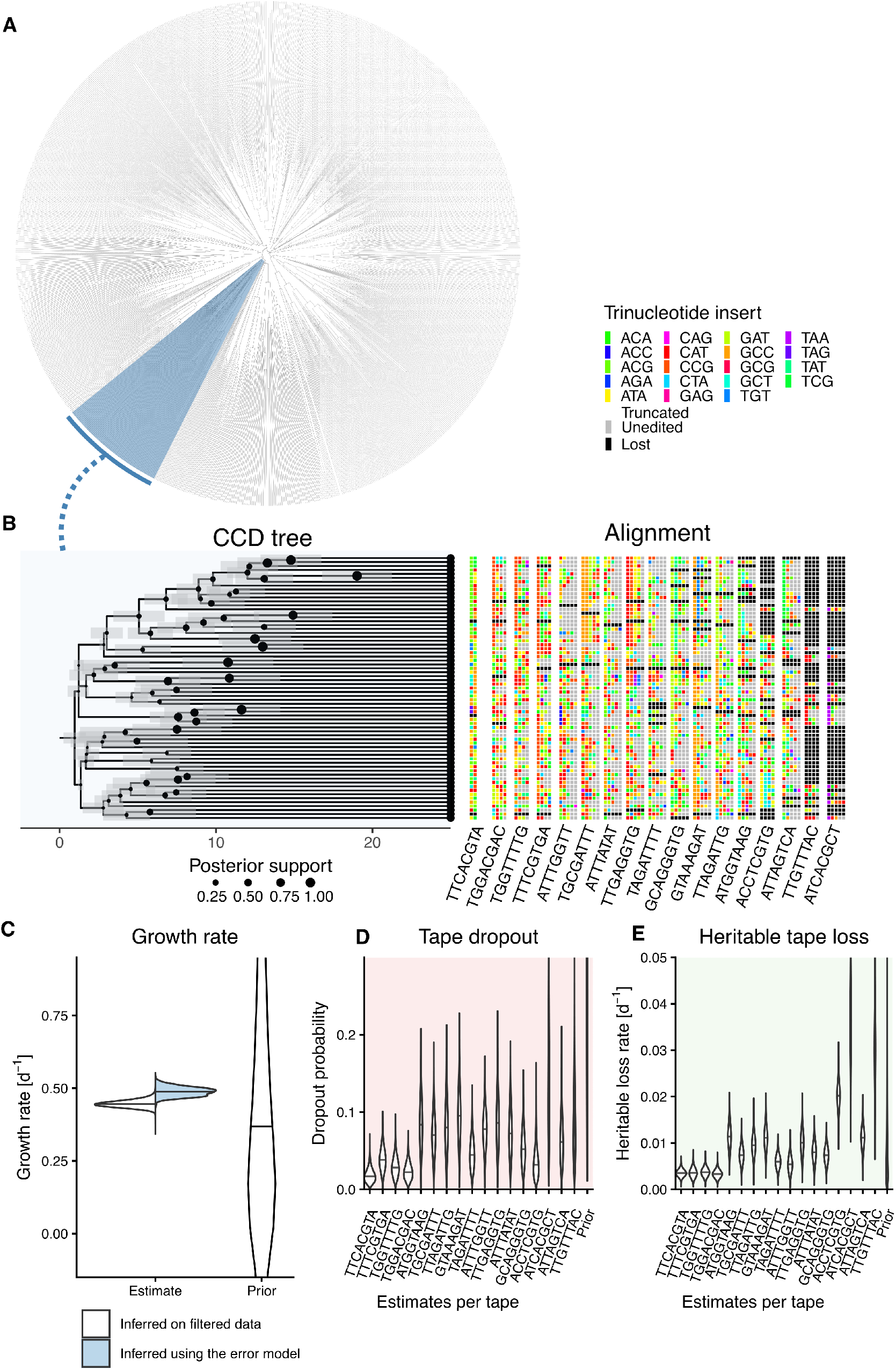
Phylogenetic estimates for a monoclonal expansion of HEK293T cells using SciPhy and our mechanistic model of barcode loss. **A** Reconstructed phylogeny: Conditional clade credibility (CCD) summary tree inferred using our mechanistic loss model on a subsample of 1,000 DNA Typewriter sequences. A representative subclade is highlighted in blue. **B** Subclade detail: Enlarged view of the annotated clade from A. Node sizes represent posterior support, and grey bars indicate the 95% HPD intervals for node heights. **C** Growth rate estimation: Comparison of posterior distributions for the net growth rate. Results are shown for the original, filtered baseline estimates reported in [16] (white) and our new mechanistic model utilizing the unfiltered dataset (blue). Posterior distributions for the parameters of our mechanistic loss model are shown for the **D** stochastic dropout probability and **E** heritable loss rate. In all violin plots (C–E), central horizontal lines denote the posterior median.

Similarly, all barcodes were estimated to have a higher editing rate using the loss model compared to the filtered data (ranging from 0.138 to 0.295 *d*^−1^, see

Appendix figure 12 A). We also report estimates for the loss model parameters in Figure 4 C and D. These exhibit considerable variation between barcodes, with posterior medians ranging between 0.003 and 0.034 *d*^−1^ for the heritable loss rate, and a tape dropout probability ranging from 1.7% to 13.5% per barcode. Although the heritable loss rate is difficult to validate independently, the inferred tape dropout probabilities match the low-dropout regime expected for this construct design, where strong promoter activity (eEF1*α*) combined with dedicated 3^*′*^ RT capture primers ensures efficient transcript recovery in single-cell 10*×* sequencing experiments (as designed in [2]). Finally, the inferred sampling proportion was slightly lower under the loss model (posterior median *≈*0.00009, Appendix Fig. 12D). This highlights how jointly estimating the sampling proportion alongside growth parameters yields a more coherent fit to independent reference measurements of the expansion dynamics than fixing it *a priori*. Although birth, death, and sampling parameters are subject to theoretical unidentifiability in an unconstrained birth-death-sampling model, fixing the experiment’s origin alongside the use of realistic priors achieves practical identifiability for the primary parameter of interest here—the net growth rate. Our re-analysis of the gastruloid dataset from [16], yields similar patterns of increased editing and growth rates (see Fig. 15). Specifically, the error model identifies a dropout probability of 1.64% and elevated growth estimates—particularly in the final expansion phase (0.595 *d*^−1^ vs. 0.529 *d*^−1^ with filtering)— while still providing dynamics that remain in qualitative agreement with reference data from [28].

### Analysis of sparse, high-entropy alignments

A crucial advantage of our mechanistic loss model is that it enables the construction of datasets that include barcode alignments selected for their information content rather than strictly requiring complete (lossless) alignments. In multi-copy single-cell lineage tracing experiments, up to *∼*100 independent barcode copies can be integrated into the genome [29]. Analyzing all barcode alignments can become computationally prohibitive, and in particular wasteful when many barcodes remain unedited, contributing virtually no phylogenetic signal. Given a fixed set of cells, traditional pipelines are forced to select barcodes purely based on sequence completeness, often discarding highly informative, heavily edited barcodes simply because their alignments contain missing sequences. In contrast, our framework supports lossy alignments, enabling barcode selection based on editing signal rather than completeness. We showcase this advantage by applying our model to previously excluded gastruloid data [16], comprising 6 biological replicates pooled pairwise across 3 sequencing lanes (2 replicates per lane). These replicates were originally deemed unsuitable because their complete barcode alignments were poorly edited and thus lacked sufficient phylogenetic signal to reconstruct high-confidence trees of an acceptable size (*>* 100 cells). To quantify the utility of sparser yet more informative barcodes, we employed Shannon entropy as a measure of phylogenetic information content (see Materials and Methods), reflecting the diversity and complexity of the recorded editing patterns. By demultiplexing replicates via barcode clustering and filtering for barcodes with Shannon entropy *>* 2, our framework successfully generated nine viable subsamples across hundreds of cells (derived from individual replicates within the 3 sequencing lanes; see Materials and Methods). Figure 5 illustrates the results of barcode selection (A), tree inference (B), and growth rate estimation (C) for the three subsets with the highest information content, resulting in reconstructed trees of significant size: Lane 3 Subsample 1 (Mean Entropy: 4.14; Tree size *N* = 462), Lane 3 Subsample 2 (Mean Entropy: 3.61; Tree size *N* = 482), and Lane 2 Subsample 1 (Mean Entropy: 3.48; Tree size *N* = 261; see Appendix Fig. 16 for all others). Overall, seven out of nine subsamples successfully recovered the expected growth dynamics: a sustained growth rate until day 7, followed by a decrease coinciding with the onset of gastruloid elongation [16, 29] (see Appendix Fig. 17). We observe that the agreement with the reference growth trajectory [28] is generally stronger in subsamples with higher mean entropy, suggesting that information content is a reliable predictor of a dataset’s capacity to resolve growth dynamics. An exception is seen in Lane 3 Subsample 3; despite a high mean entropy (3.25), a lower cell recovery (mean *N* = 105, or 47.1% of the 223 available cells) compared to other Lane 3 subsets likely introduced sampling noise that obscures the expected late-stage growth decrease.

**Figure 5.**
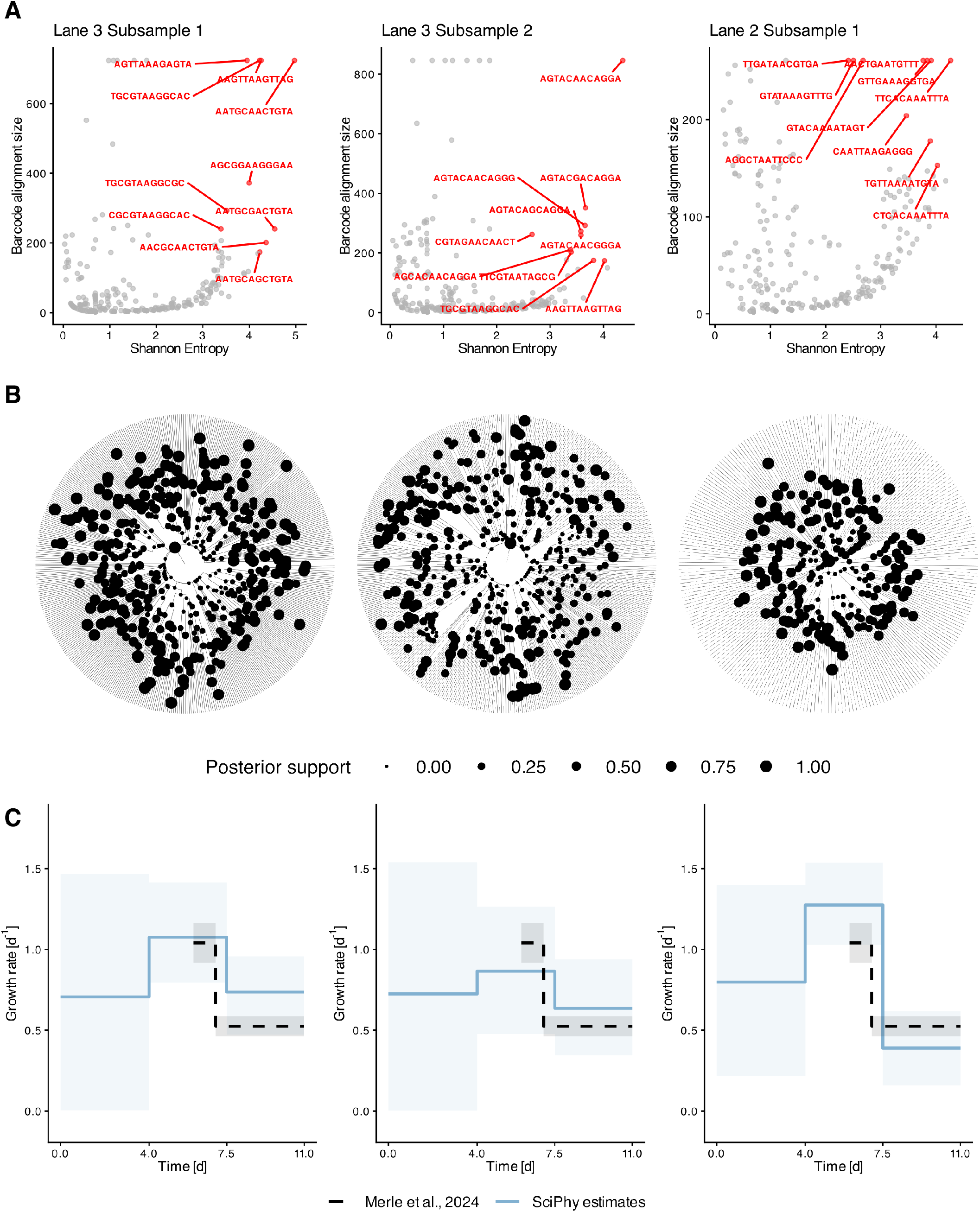
High-information alignment construction and growth rate estimation in murine gastruloids. **A** Targeted alignment selection: Relationship between Alignment Entropy (Shannon entropy) and alignment size for 3 barcode subsets extracted from monoclonal murine gastruloids. Our mechanistic loss model enables the selection of high-entropy, sparse barcodes that are typically discarded; here, we include up to 10 barcodes with maximum entropy present in at least 20% of cells. **B** Phylogeny reconstruction: Representative CCD trees inferred for each selected barcode dataset, demonstrating consistent topology recovery despite varying levels of barcode loss. **C** Growth rate estimation: Posterior estimates of the net growth rate through time (median and 95% HPD; blue) across datasets. Estimates are compared against reference values from [28] (grey/dashed), demonstrating that our model accurately recovers biological growth dynamics from previously inaccessible, lossy data.

**Figure 6.**
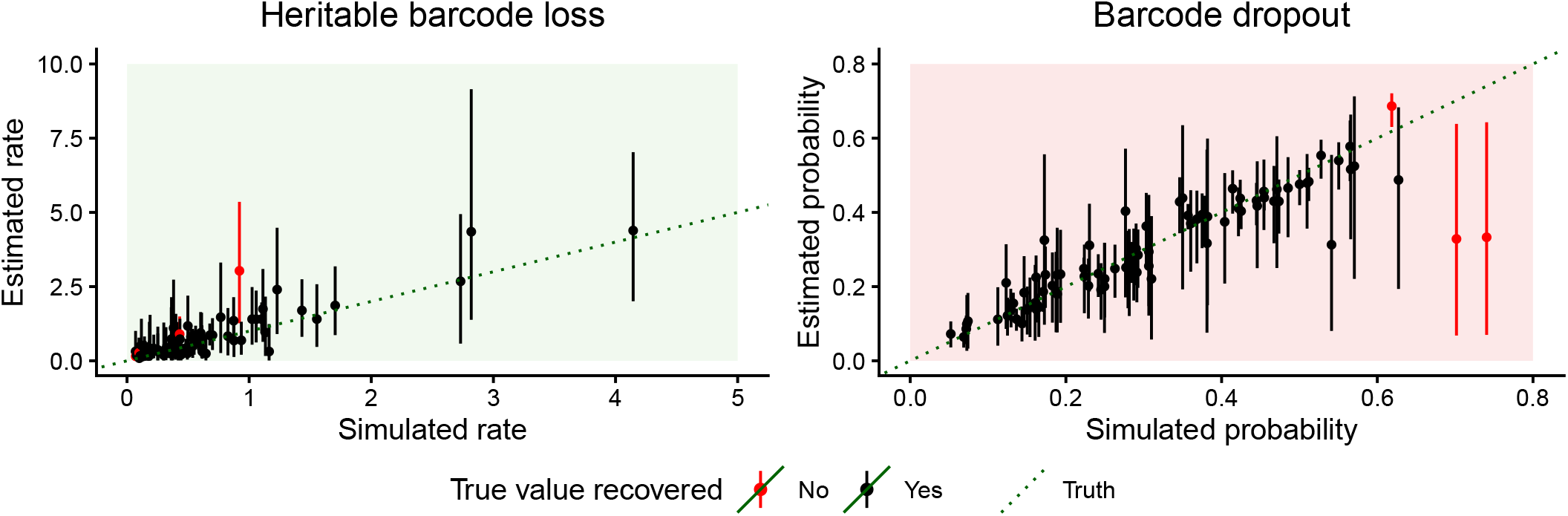
In-silico validation of the TiDeTree model extension. Validation results for theheritable loss rate (left), and dropout probability (right). Dots represent posterior medians and lines the 95% Highest Posterior Density (HPD) intervals, against a line (green) indicating the identity (*y* = *x*) baseline of a perfect estimator. Points and intervals are colored black if the 95% HPD interval contains the true value and red otherwise. The model achieves coverage of 93%, 96%, and 94% for the editing rate, heritable loss rate, and dropout probability, respectively.

**Figure 7.**
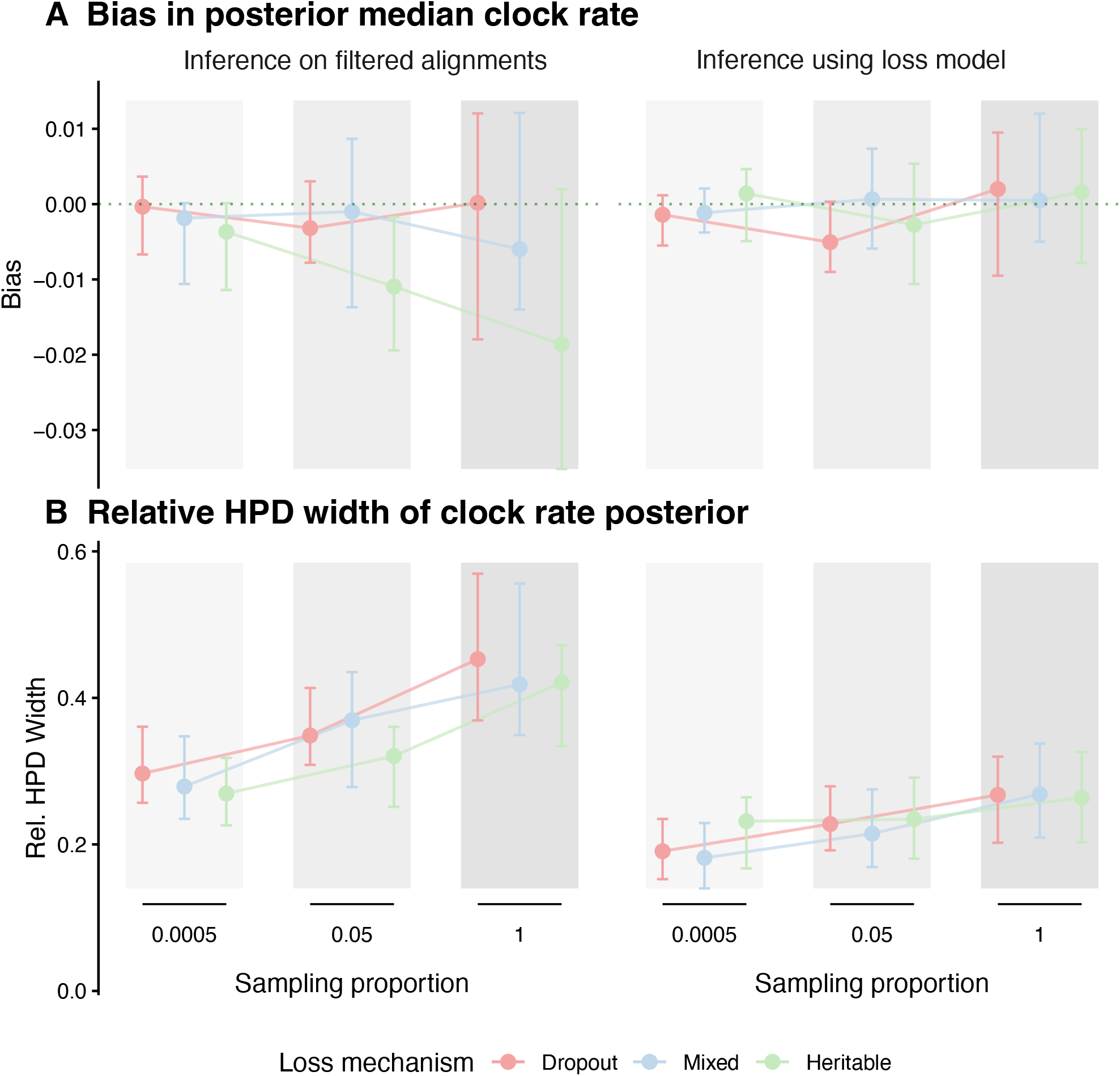
Simulation-based comparison of editing (clock) rate estimation under data filtering and mechanistic loss modeling. **A** Absolute bias and **B** Mean relative HPD width The estimation is compared varying loss scenarios (colors) and tree shapes (modulated by sampling proportion, background shading). Points represent the mean performance across replicates, with error bars denoting the 95% interval of the estimates. In A, the horizontal dotted line indicates an ideal bias of zero.

**Figure 8.**
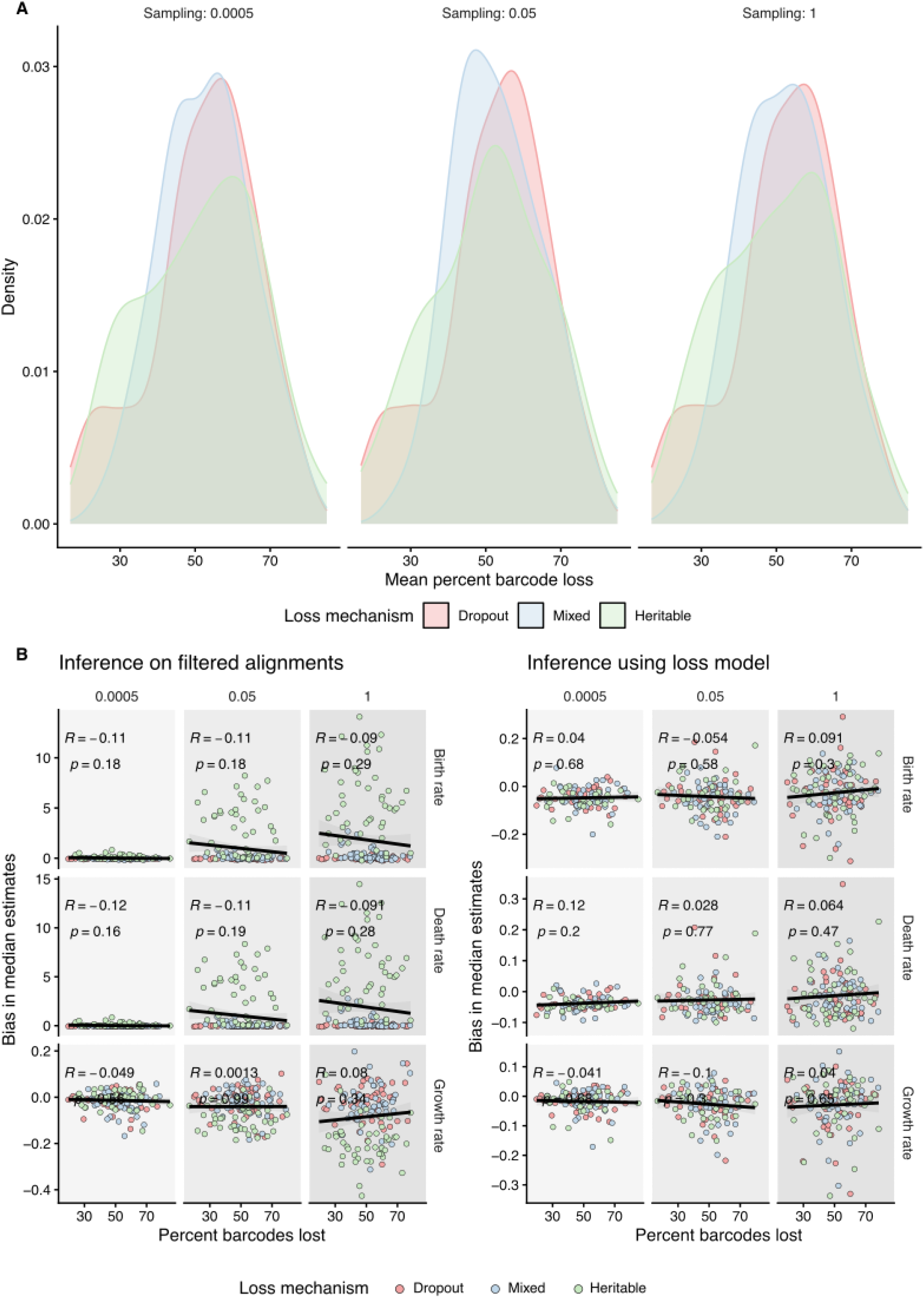
Impact of overall barcode loss on parameter bias. **A** Density distributions of the mean percentage of barcode loss over all simulated datasets across sampling proportions (columns) and loss mechanisms (colors). Total data loss remains consistent at approximately 50% across all scenarios, isolating the effect of the loss mechanism rather than the loss magnitude. **B** Correlation between the percentage of data loss and bias in birth (*λ*), death (*µ*), and net growth (*r*) rates.

**Figure 9.**
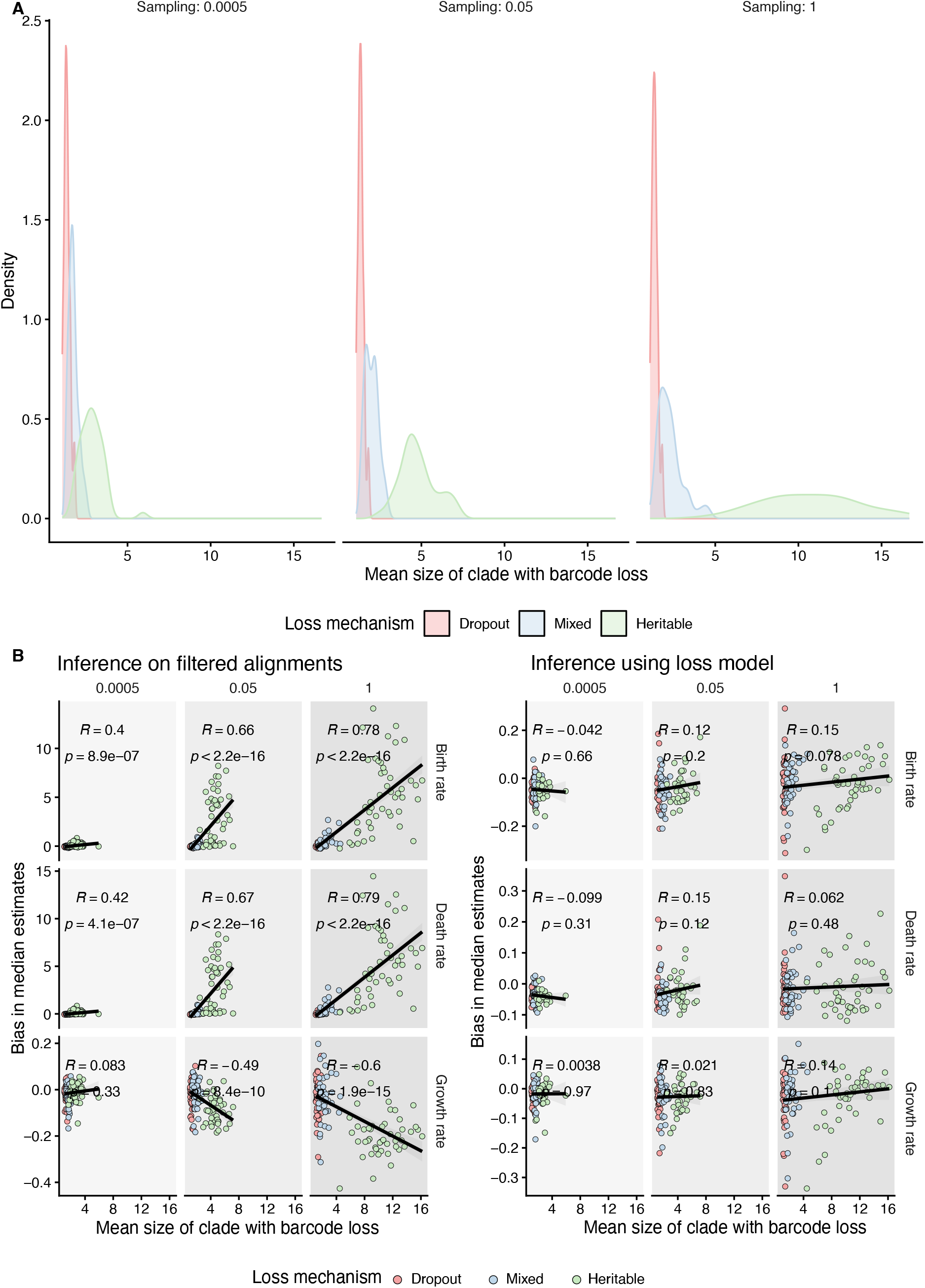
Impact of lost clade size on parameter bias. **A** Density distributions showing the mean size of clades affected by data loss over all simulated datasets across the three sampling proportions (columns) and three loss mechanisms (colors). Heritable loss (green) results in larger monophyletic clades with lost barcodes compared to dropout (pink), a distinction that intensifies as sampling density increases. B Correlation between lost clade size and bias in birth, death, and net growth, rates.

**Figure 10.**
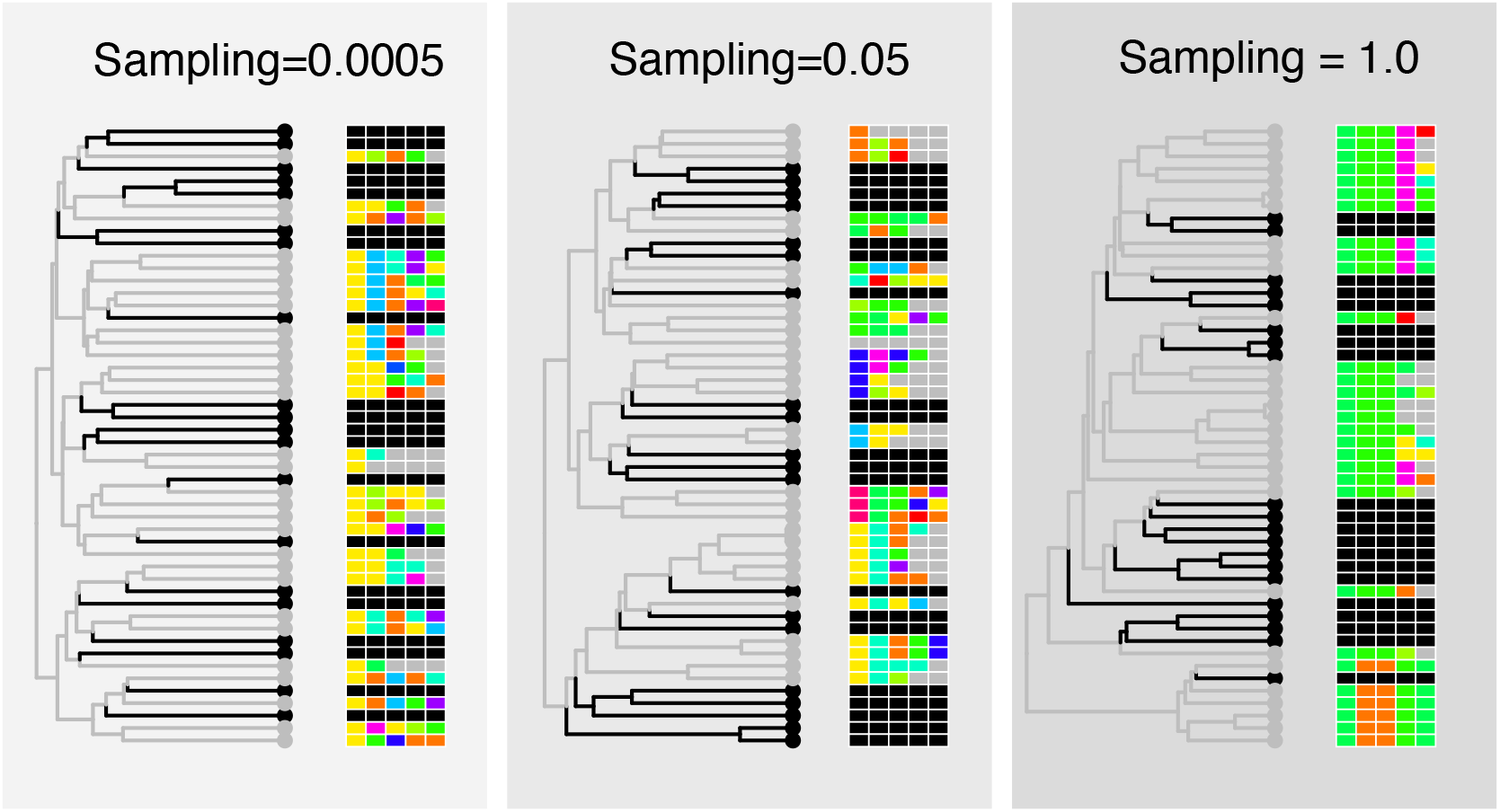
Explanatory schematic illustrating the effect of sampling proportion on tree branching and heritable barcode loss patterns. Holding total tree size and heritable loss rate constant, increasing the sampling proportion (left to right) shifts branching toward the root. This topological shift causes heritable loss events to affect larger, highly clustered clades, resulting in increasingly aggregated missing data across the alignment.

**Figure 11.**
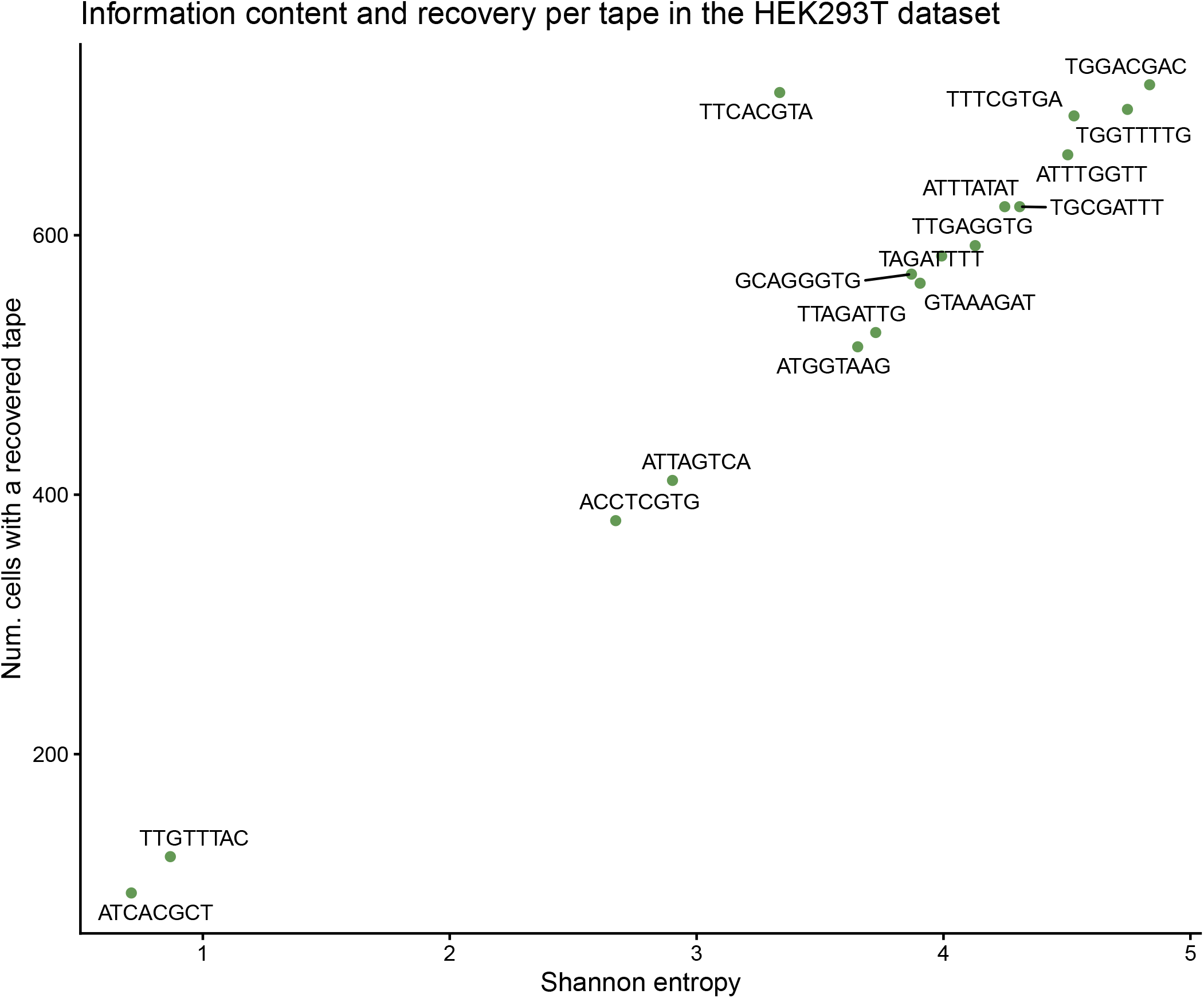
Information content and barcode recovery per tape in the HEK293T dataset. Scatter plot showing the relationship between Shannon entropy (x-axis) and cell recovery (y-axis) for each target barcode or “tape” used in the dataset.

**Figure 12.**
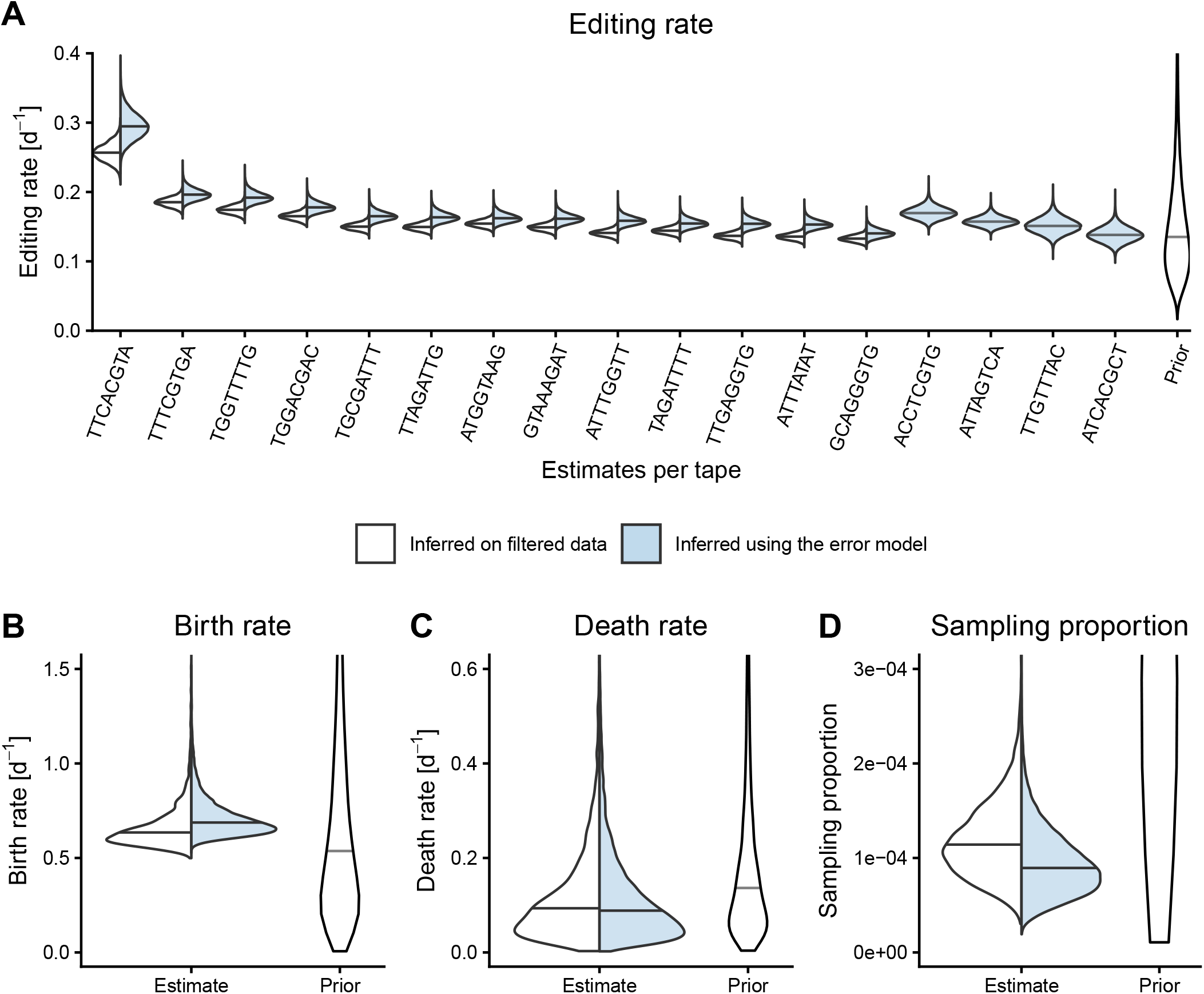
Comparison of phylogenetic estimates obtained in SciPhy for the monoclonal growth of HEK293T cells harboring DNA Typewriter barcodes, when filtering out lossy cells or with our model extension co-estimating the sampling proportion. We show estimates of the editing rate per barcode (**A**) and birth and death rates (**B** and **C**) estimated using both approaches.

**Figure 13.**
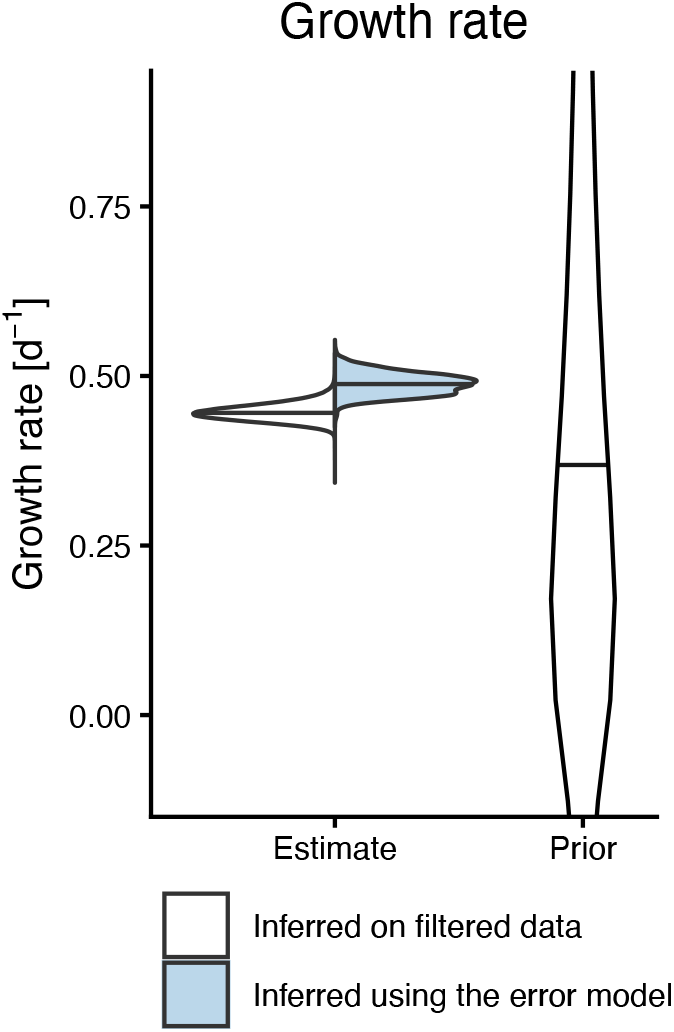
Comparison of estimates of the growth rate obtained in SciPhy for the monoclonal growth of HEK293T cells harboring DNA Typewriter barcodes, when filtering out lossy cells or with our model extension, with the sampling proportion fixed to 0.0008.

**Figure 14.**
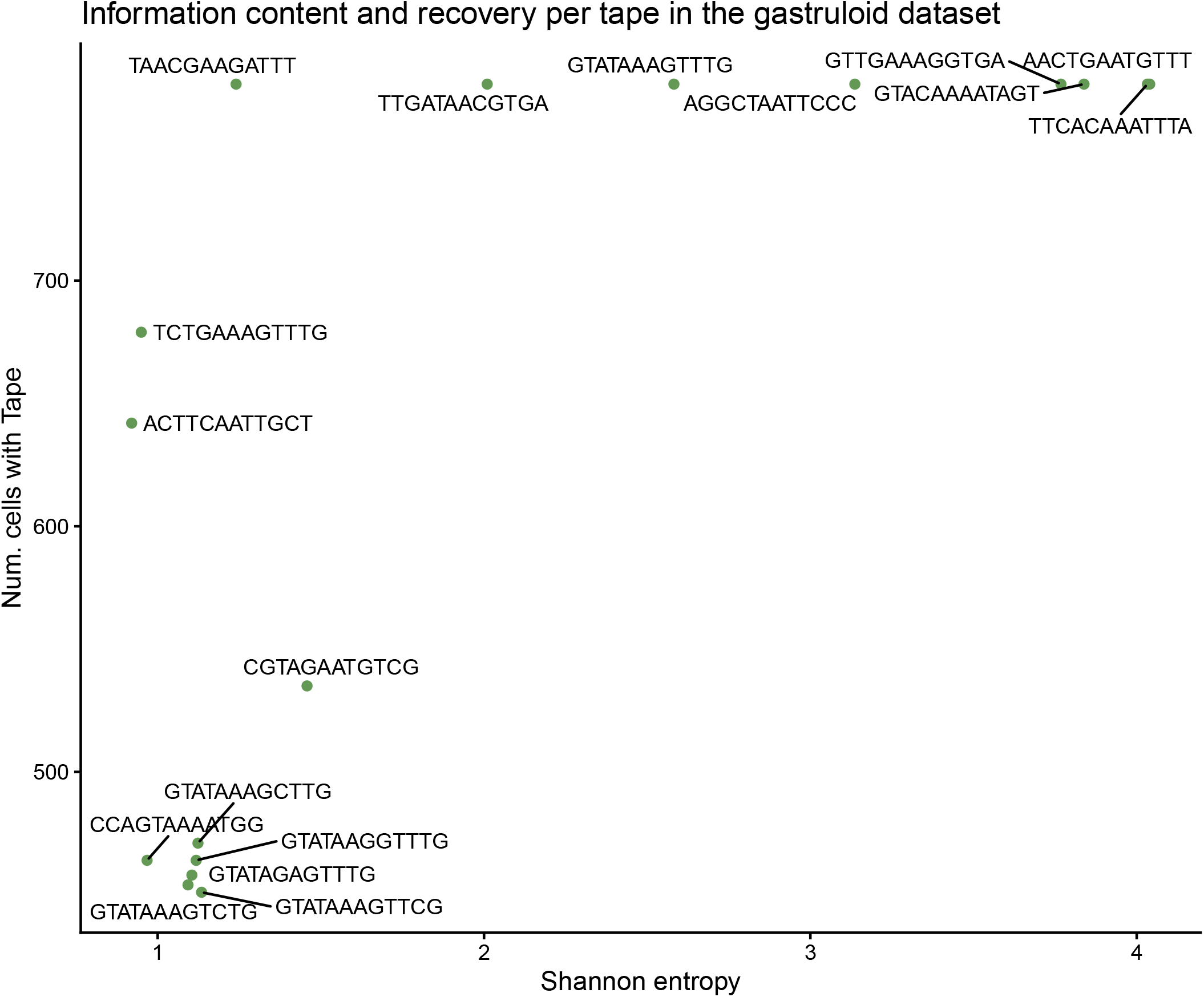
Information content and barcode recovery per tape in the gastruloid dataset analysed in [16]. Scatter plot showing the relationship between Shannon entropy (x-axis) and cell recovery (y-axis) for each target barcode or “tape” used in the dataset.

**Figure 15.**
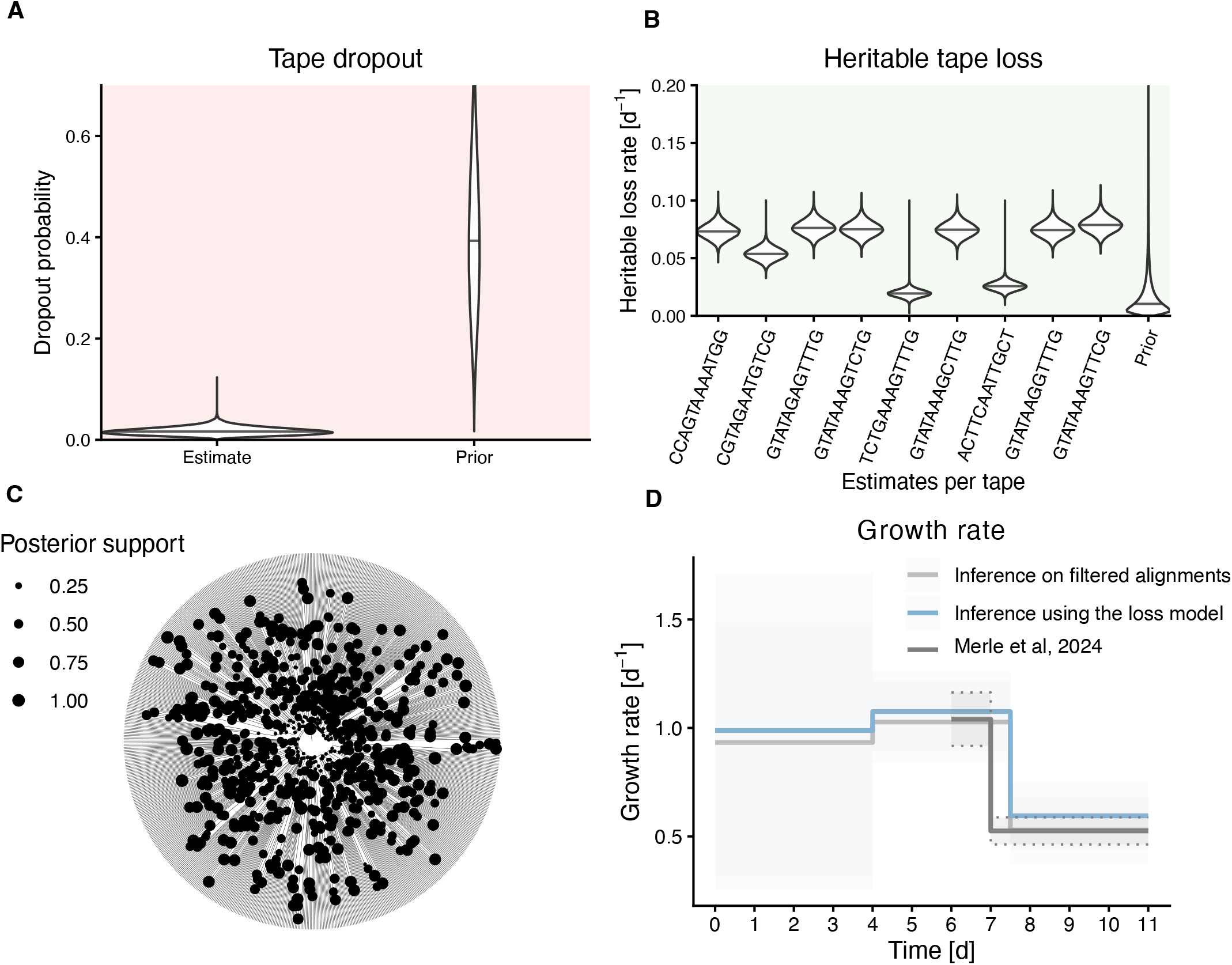
Comparison of phylogenetic estimates obtained in SciPhy for the growth of mouse gastruloid from a founding mouse embryonic stem cell (mESC) cell harboring DNA Typewriter barcodes, when filtering out lossy cells or with our model extension. In **A** and **B** we show estimates of the editing rate per barcode and growth rate estimated using both approaches. In **C** and **D** we show estimates of the heritable loss rate per barcode, and dropout probability using the loss model.

**Figure 16.**
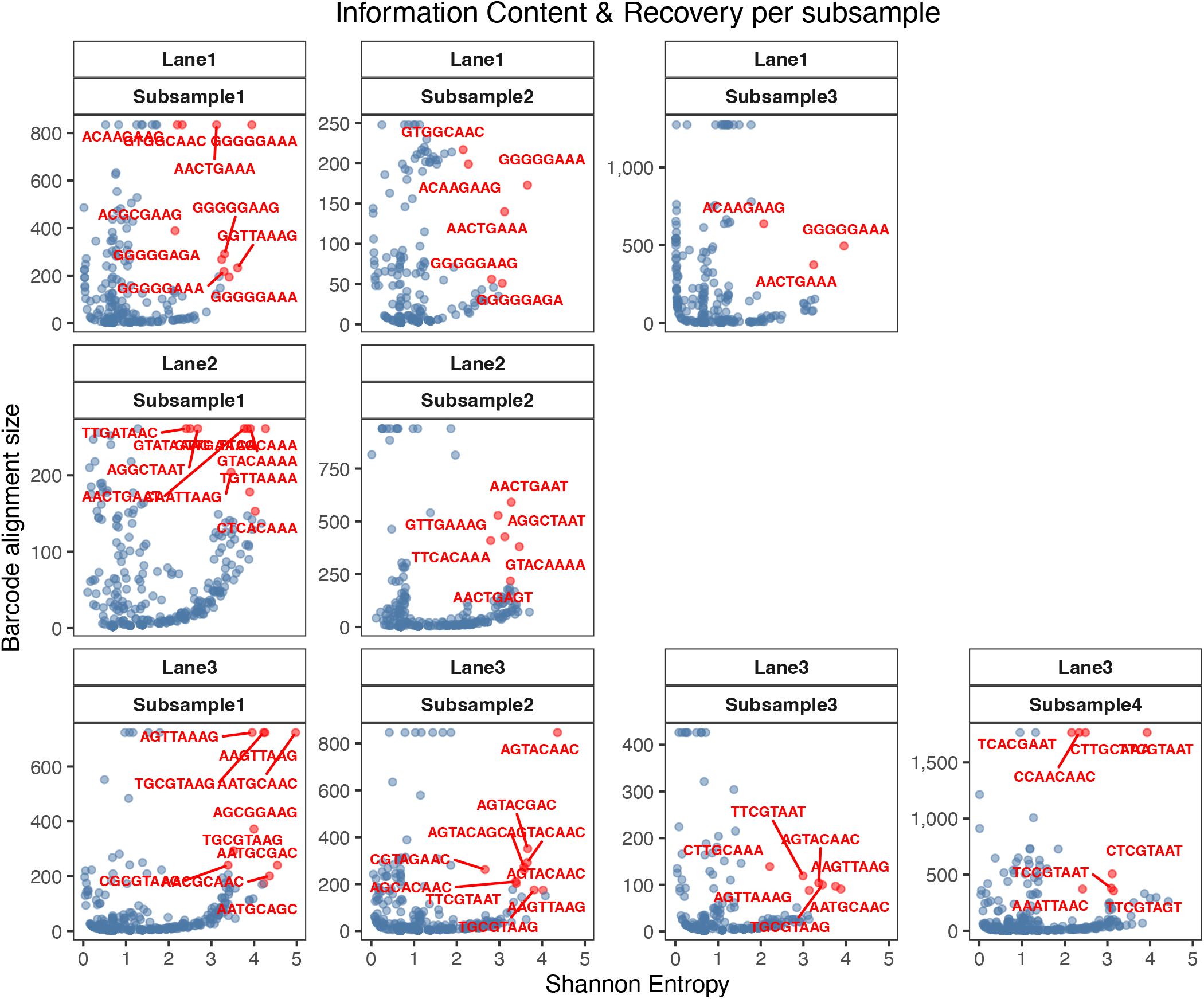
Information content and barcode recovery across subsamples of gastruloid replicates. Scatter plot shows the relationship between Shannon entropy (*x*-axis) and total cell recovery (*y*-axis) for every unique target barcode (“tape”) across all subsamples. Red highlights indicate specific barcodes included for analysis (with Shannon entropy ¿ 2 and recovery in more than 20% of cells) with labels corresponding to their identifiers. Facets are organized by experimental lane and subsample. The subsamples included in main are the top 3 datasets by mean alignment entropy, Lane 2 and 1 subsamples 1 and 2 and Lane 3 subsample 1

**Figure 17.**
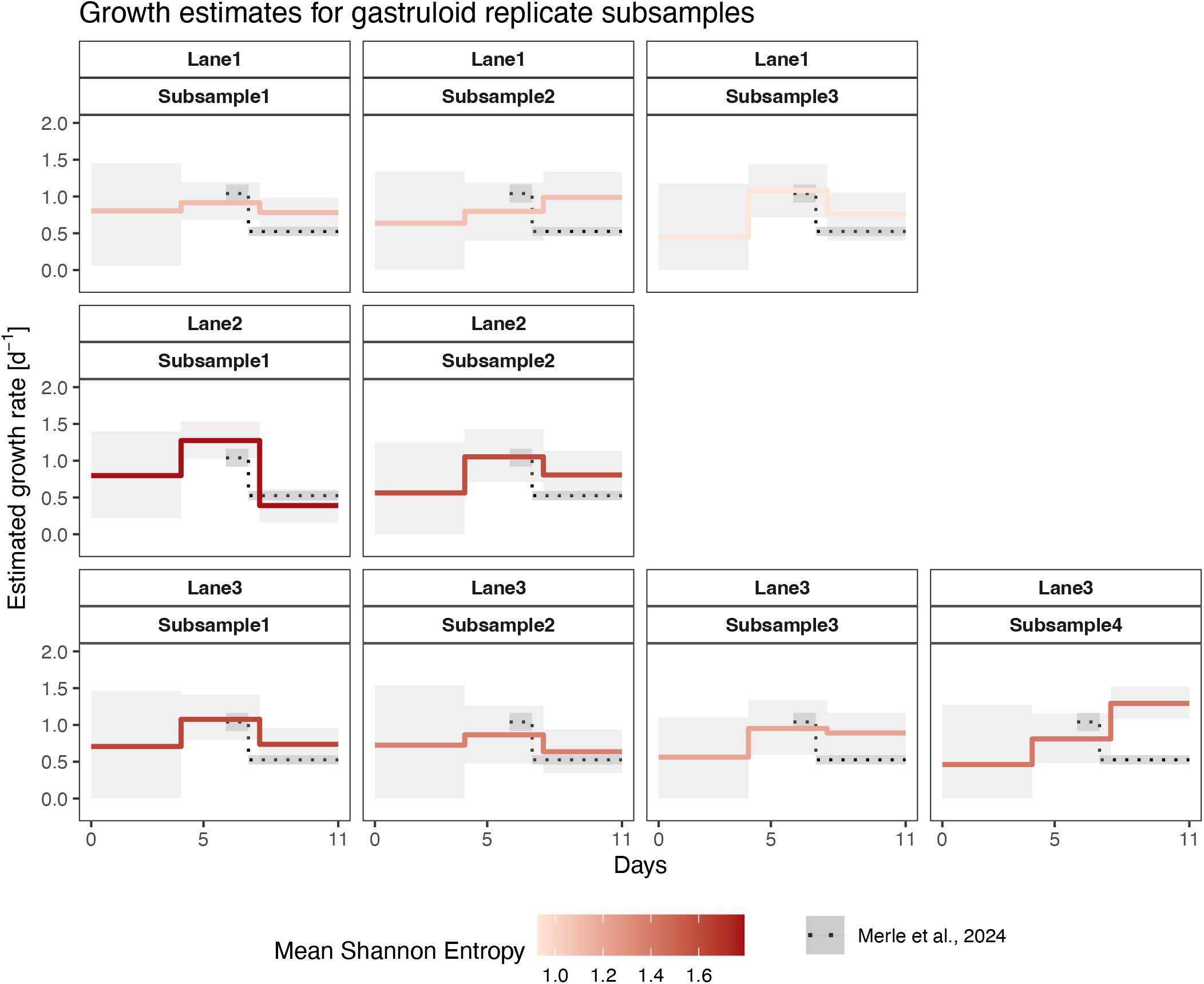
Reconstructed growth dynamics across subsamples of gastruloid replicates. Inferred growth rates obtained using the SciPhy framework, incorporating a barcode loss model under a birth-death-sampling process. Data represent subsampled populations derived from replicates of gastruloids derived from mouse embryonic stem cells (mESCs) harboring DNA Typewriter barcodes. Each panel displays the median posterior growth rate (solid colored lines) and 95% HPD intervals (grey ribbons), with lines colored according to the mean Shannon entropy of the corresponding dataset. The black dotted line and shaded region represent reference growth data from Merle et al. (2024). The subsamples included in main are the top 3 datasets by mean alignment entropy, Lane 2 and 1 subsamples 1 and 2 and Lane 3 subsample 1.

## 4 Discussion

In this study, we demonstrate that accounting for mechanisms of barcode loss can be essential for accurate phylogenetic and phylodynamic inference from lineage tracing alignments. While aligning with previous maximum likelihood work showing that explicit modeling of heritable silencing and dropout improves tree reconstruction accuracy [12], the key novelty of our study lies in the use of a Bayesian implementation to examine how unmodeled data loss distorts jointly inferred population growth parameters in phylodynamic analyses. Crucially, our simulation results showed that by circumventing the ad-hoc practice of discarding incomplete data, our implementation shields cell population growth dynamic parameters, such as cell division and death rates, from systematic biases. Beyond correcting these biases, our framework offers a major practical improvement by enabling researchers to analyze all available lineage tracing data, regardless of its level of sparsity or how those missing data are distributed across cells and barcodes.

We validated these simulation and practical insights through two empirical applications. First, in our phylodynamic analysis of a monoclonal expansion of immortalized cells (HEK293T) in culture, the inclusion of all available barcodes—including those previously filtered out—yielded a higher growth rate estimate than previously inferred in [16]. This estimate, corresponding to a 28-hour doubling time, matches the known biological ground truth for this cell line and corroborates our simulation finding that ignoring missing data can underestimate growth.

Second, in our phylodynamic analysis of a murine gastruloid dataset, we showcased the utility of the loss model in rescuing replicates previously deemed too sparse for inference. By using Shannon entropy as a proxy for phylogenetic information content, we were able to systematically select subsets of cells and barcodes that traditional filtering workflows had excluded. Our results indicated that 7 out of 9 such subsamples successfully recovered the expected developmental growth dynamics, capturing a decrease in growth that coincides with gastruloid elongation. Together, these findings demonstrate that the diversity of editing patterns (entropy) is a critical driver of reconstructive accuracy, potentially more so than the absolute completeness of alignments.

The observed effects of our loss model on tree reconstruction echo broader findings in single-cell error modeling. As noted by [18], failing to account for noise can lead to an artificial inflation of total tree length. Consistent with this, our simulations demonstrated that that estimated time-scaled phylogenies are improved by our model; by attributing missing data to heritable silencing or dropout, our framework provides a more accurate calibration of the tree branches. However, the impact of these error models likely extends beyond total tree length to the finer topological structure of the tree. Following the approach in [18], future work should investigate how accounting for data sparsity influences specific branching patterns. For example, evaluating whether our loss model affects the resolution of terminal splitting events or alters tree balance could clarify specifically how systematic barcode loss might distort inferred phylogenies.

Our framework has several limitations. First, it does not account for potential sources of noise beyond data loss, such as barcode editing errors or consensus sequence uncertainty in RNA-based protocols. Although these error rates are generally thought to be low (*≤*1%), incorporating them into future extensions may further improve topological and parameter accuracy.

Second, a discrepancy appears when comparing our simulation results with empirical estimates under low sampling intensity. While our simulations suggested that phylodynamic inference under low sampling proportions is generally robust to data-filtering biases, our HEK293T analysis revealed a significant correction in growth estimates—shifting the inferred doubling time to 28.0 hours—even at an extremely low sampling proportion (*ρ ≤* 0.0008). One hypothesis for this discrepancy is that biological barcode loss may not be strictly uniform across lineages. Because our simulation study pinpointed the deletion of large, structured clades as the primary driver of bias, the manifestation of this bias in empirical data may imply that loss is exacerbated within specific lineages—perhaps arising from localized epigenetic silencing or early progenitor dropouts that create structured gaps. Reassuringly, our model’s ability to rescue the true growth rate even in this context highlights its robustness to such potential biological heterogeneities, which could be formally modeled in future clade- or time-dependent extensions [30].

Finally, while there is consensus across the literature that missing data generally should not be ignored, an alternative approach is to bypass explicit mechanistic modeling. For instance, recent maximum-likelihood work [31] demonstrated that missing data can be treated as an “ignorable” process without biasing branch length estimates. Extending this non-mechanistic, computationally efficient paradigm to a joint Bayesian framework represents a promising future direction. Beyond SciPhy and TiDeTree, such strategies can serve as a scalable alternative for modeling loss in GESTALT-like data [13], which suffer from more complex, within-barcode types of loss. Adapting these simplified missing-data formulations into BEAST 2, or combining them with our loss framework, provides a potential path to handle such loss patterns efficiently, bypassing the heavy computational overhead associated with likelihood calculations in GAPML [10] and GABI [11].

In summary, we provide a loss model for single-cell lineage tracing that is fully embedded within the BEAST 2 software ecosystem, scalable to datasets of up to approximately 1,000 cells—the typical upper bound for joint Bayesian phylogenetic inference tools. This framework enables researchers to fully leverage the high-entropy, high-sparsity lineage tracing datasets currently being generated at scale. Because our framework is built as a modular platform that facilitates further modeling extensions and computational optimization, we envision this implementation becoming increasingly relevant as next-generation datasets combine lineage tracing with additional single-cell modalities to interrogate biological processes of growing complexity.

## 5 Materials and Methods

### Model of loss for lineage tracing barcodes

We describe a model for the processes underlying the loss of lineage tracing genomic constructs, hereafter referred to as “barcodes”. These barcodes consist of genomic loci that accumulate permanent and irreversible edits through various genome-editing modalities [4, 13, 29]. In the following, we consider a general experimental setup in which a founding cell harboring several barcodes undergoes proliferation, during which barcodes may accumulate changes, and after which the resulting population is sampled for single-cell sequencing. The resulting single-cell readouts are usually incomplete, missing one or several of the original barcodes. We model this sparsity as the result of two distinct processes: heritable loss, where barcodes are rendered inaccessible in a stable manner during proliferation (e.g., via transgene silencing or deletions), and stochastic dropout occurring during the single-cell sequencing workflow.

We consider the following notations. A single founding cell at time 0 carries *k* barcodes *b*_1_, …, *b*_*k*_ with each *N* target sites, and undergoes proliferation for an experimental duration of *t*_*e*_. We denote the complete set of *k* barcodes as *B*, where:

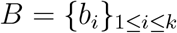

At any time *t* during the experiment, each descendant *m* of the founding cell carries a set of at most *k* of the original barcodes, denoted as *S*_*m,t*_, where *S*_*m*_ *⊆ B*. A lossy set of barcode sequences is defined as any set *S*_*m,t*_ containing less than the full set of *k* barcodes, i.e. |*S*_*m,t*_| *< k*.

We define two potential sources of loss for these barcodes:

1. Heritable loss: barcodes are irreversibly lost at rate *r*_*l*_ during the lifetime of a cell. A barcode that is lost cannot be reacquired by descendant cells. This barcode loss corresponds to a Poisson process with a rate *r*_*l*_ throughout the lifetime of the cell.
2. Dropout: upon sampling and sequencing, a barcode is lost with probability *p*_*d*_. Barcode dropout is therefore the result of a single Bernoulli trial with probability *p*_*d*_ at the end of the experiment.
3. Each barcode *b*_*i*_ may have its own independent loss rate and probability, *r*_*l,i*_ and *p*_*d,i*_, for instance due to effects of the genomic context of a barcode.

### Probability of barcode loss

Given the processes described above, we distinguish two cases to calculate the probability that a given barcode *b*_*i*_ is lost in any time interval (*t*_*a*_, *t*_*b*_) along a cell lineage. In case 1, the time interval does not lead to the end of the experiment *t*_*b*_ *< t*_*e*_. In this case, a barcode can only be missing as a result of a heritable loss during the time interval, with probability:

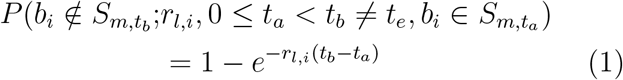

In case 2, the time interval leads to the end of the experiment and sequencing, *t*_*b*_ = *t*_*e*_. There, barcode loss can result from either heritable loss during the time interval itself or sequencing dropout at its end. In such a case, the corresponding probability that barcode *b*_*i*_ is lost is:

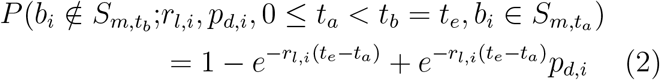

The probability that any given barcode is lost over the time interval from the start to the end of the experiment (i.e., substituting *t*_*a*_ = 0, in case 2) is:

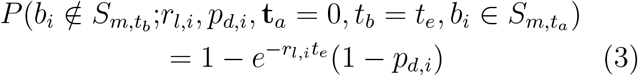

### Tree likelihood

The components described above extend the models from [9, 16], allowing to evaluate the probability of a lossy dataset *D*_*l*_ given a cell phylogeny *T*, loss parameters (*r*_*l*_, *p*_*l*_), and editing parameters *η*, denoted by *P* (*D*_*l*_; *η, r*_*l*_, *p*_*l*_, *T* ). This is the likelihood of the parameters given the alignment.

This likelihood is computed using Felsenstein’s pruning algorithm [32], which marginalizes over the ancestral states of each barcode. To handle loss across different barcode designs, our approach adapts to how marginalization is defined in each model’s existing likelihood implementation:

- **In SciPhy (Sequential):** Because SciPhy’s underlying model marginalizes over states that represent entire barcode sequences, we are required to utilize a “wildcard” state to avoid explicitly summing over this high-dimensional space. We thus augment the barcode state space with a “silenced” state (heritably lost barcodes), a “dropout” state (stochastically lost at tips), and a “wildcard” state representing the set of all possible sequences that are not lost. For tips with missing barcode sequences, the likelihood is then partitioned between the dropout and silenced states; the wildcard state is assigned zero likelihood, as observing a missing sequence excludes non-lost states at that tip. Because internal node states are formed by intersecting descendant state sets (excluding tip-only dropout states), the wildcard can propagate through internal nodes to represent unobserved (and non-silenced) sequence states (Fig. 18A).
- **In TiDeTree (Unordered):** Because TiDeTree natively marginalizes at the level of individual, independent sites within barcode sequences, a wild-card state is unnecessary. Instead, missing observations are handled by setting the tip likelihoods to *p*_dropout_ across all potential non-silenced site states. This effectively treats dropout as an uninformative observation at the tips, while the “silenced” state is directly integrated as an additional state in the transition matrix (Fig. 18B).

**Figure 18.**
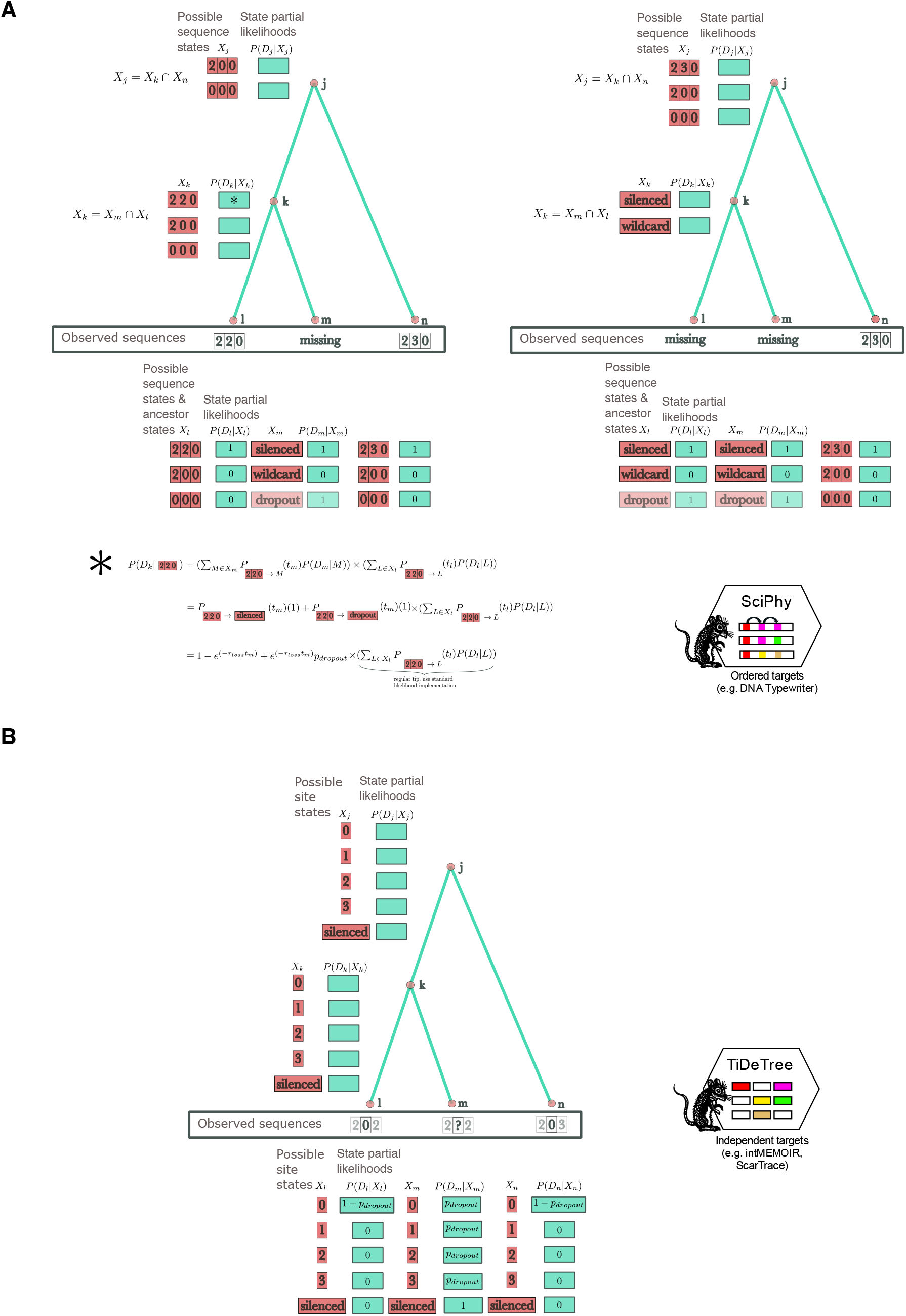
Integrating the loss model into the likelihood implementations of SciPhy and TiDeTree. **A** SciPhy: Partial likelihoods are recursively calculated for sets of barcode sequence states, determined at each node as the intersection of descendant state sets (excluding the dropout state, which can only appear at tips). We extended this framework to accommodate missing data by introducing a dropout state, a silenced state, and a wildcard state (representing any valid barcode sequence state that was neither silenced nor dropped out). Two examples illustrate how missing barcode sequences dictate ancestral states at internal nodes, alongside an example of the resulting partial likelihood calculation. **B** TiDeTree: Conversely, TiDeTree calculates partial likelihoods for individual sites within the barcodes, allowing it to operate on a fixed state space of all possible characters at each node—which already included a silenced state. Consequently, the TiDeTree implementation is extended by directly factoring the dropout probability into the site-specific partial likelihoods at the tips, as shown in the example.

Using these expanded state spaces, we apply the recursions described in [16] and [9] to compute partial likelihoods at internal nodes.

Integrating the loss model into the SciPhy framework is achieved at a negligible, constant computational cost. For a set of *k* barcodes of length *N*, augmenting the framework with the “silenced”, “dropout”, and “wild-card” states yields a theoretical maximum state space of size *N* +3. This bounds the maximum potential state transitions to (*N* +3) *×* (*N* +3), a minimal increase from the original (*N* + 1) *×* (*N* + 1) space. Specifically, the “silenced” state increases the number of potential internal node transitions by one, and the “wildcard” state prevents a combinatorial explosion of the state space by collapsing transitions across unobserved edited states. Additionally, as illustrated in Fig. 18, these augmented states only propagate within subtrees composed exclusively of missing sequences. Consequently, for a dataset of *m* cells, the total likelihood computation remains tightly bounded, scaling polynomially with barcode length and strictly linearly with the number of cells, maintaining a computational complexity of *O*(*kmN* ^2^).

The TiDeTree implementation similarly scales efficiently with the augmented state space. Because it already incorporates a single “silenced” state into its transition matrix, handling stochastic dropout requires only a simple modification of the initial tip likelihoods, demanding no alterations to the core pruning algorithm.

### Implementation

The barcode loss model is integrated into the SciPhy and TiDeTree packages for the BEAST 2 phylogenetic platform, enabling both the simulation of datasets under these models and Bayesian inference under a unified framework. For simulation, we extended the following classes: SimulatedSciPhyAlignmentHeritableMissingBcodes (SciPhy) and SimulatedAlignment (TiDeTree). These classes utilize the FEAST package [33] to generate barcode sequences along a given phylogeny, conditioned on user-specified heritable loss rates (*r*_*l,i*_) and dropout probabilities (*p*_*d,i*_). To account for data loss during inference, we extended the substitution models and likelihood components of both packages. In SciPhy, this includes the SciPhySubstitution-Model and SciPhyTreeLikelihood; in TiDeTree, the EditAndSilencingModel and TreeLikelihoodWithEd-itWindow were similarly adapted. These classes are designed to process missing barcodes, represented as *N* -tuples of “?” symbols in the input alignment, by marginalizing over missing information (using representative (wilcard/silenced/dropout) states in SciPhy or uninformative tip likelihoods in TiDeTree) within the phylogenetic likelihood calculation. The posterior distribution of the phylogenetic tree and model parameters, given lossy barcode alignments, is estimated using Markov Chain Monte Carlo (MCMC). All model specifications, including the selection of loss mechanisms and the definition of parameter priors, are configurable via XML scripts, ensuring accessibility and reproducibility within a standard BEAST 2 workflow.

### Validation and comparison to complete data

We validated the implementation and integration of the barcode loss model within the SciPhy and TiDetree packages using a well-calibrated simulation study [22]. We simulated 100 phylogenetic trees under fixed birth-death dynamics (*λ* = 0.8, *µ* = 0.2, [34] with the simulator implemented in [35] (Tab. 4). Along each tree, we generated 20 independent barcode alignments with varying degrees of data loss. Heritable loss rates and dropout probabilities were drawn from the distributions specified in Table 4. The parameter distributions were chosen to produce a broad spectrum of data loss, with the average loss per dataset ranging from 17% to 85% (also see Appendix Fig. 8A, sampling = 0.05, “Mixed”), and a mean loss of *∼*50% over all datasets. For performance comparison, we generated two additional sets of alignments: 10 complete barcode alignments (with loss and dropout set to 0.0) and a subsample of 10 barcode alignments drawn from the original 20 lossy replicates.

### Extended simulation study

To evaluate the performance of our model for phylodynamic inference, we compared it against the current standard approach: filtering out lossy barcodes and cells to generate datasets with complete alignments. This filtering process ensures that input data are compatible with traditional phylogenetic reconstruction methods and is generally implemented in three stages:

- **Target ranking:** All barcodes are ranked according to their recovery frequency.
- **Barcode selection:** A subset of barcodes is assembled iteratively to maximize barcode retention while satisfying a minimum taxon threshold, based on ranking the *i* most frequent barcodes. In our simulation study, we required that the final filtered subset of cells (defined as those containing all selected barcodes) represented at least 5% of the initial simulated population.
- **Cell selection:** Once the optimal set of barcodes is determined, any cells missing one or more of the selected targets are discarded to produce a complete alignment.
- **Effective sampling proportion calculation:** For each filtered dataset, we calculated an effective sampling proportion, defined as the product of the original simulation sampling proportion and the proportion of taxa retained after filtering. This *ρ*_*eff*_ was used to parameterize the birth-death-sampling models during inference.

To ensure a fair comparison, we adopted a conservative benchmarking strategy by evaluating an identical number of taxa across both methods. Although our mechanistic model is capable of utilizing full, unfiltered alignments (provided runtimes remain manageable), we intentionally subsampled these datasets to match the exact cell count retained by the filtering process. This constraint ensures that any observed improvements in parameter accuracy or precision are strictly attributable to our model’s explicit accounting of loss mechanisms, rather than a sample-size advantage from analyzing more tips.

We repeated the simulation and inference process for each approach across nine distinct conditions, using 50 simulated trees per condition. These scenarios comprised three sampling regimes, each paired with three barcode loss mechanisms: stochastic dropout only, a mixed scenario (dropout and heritable loss with priors centered equally on both), and heritable loss only. The prior distributions for each loss condition were chosen to ensure a consistent mean total data loss of approximately 50% across all simulations (see Fig. 8A).

The three sampling regimes were defined as follows:

- **Low sampling:** Long experimental duration (25 days) paired with a low sampling proportion of *ρ* = 0.0005.
- **Intermediate sampling:** Intermediate duration (17.3 days) paired with *ρ* = 0.05.
- **Complete sampling:** Short duration (12.3 days) with full sampling of all extant cells (*ρ* = 1.0).

The experimental duration (tree origin) was adjusted for each regime to maintain consistent final tree sizes (number of extant taxa) across conditions while achieving different branching patterns. All simulation parameter values and prior distributions are detailed in Tables 3, 4, and 6.

Performance for parameter estimation was evaluated using two primary metrics:

- **Absolute Bias:** The difference between the posterior median estimate and the true parameter value:

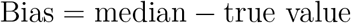
- **Precision:** The relative 95% Highest Posterior Density (HPD) interval width, normalized by the true value:

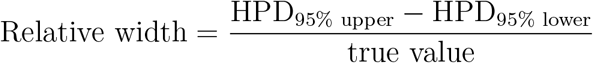

Furthermore, we quantified the accuracy of the inferred phylogenies by calculating the normalized distance to the true simulated subtrees using the Phylogenetic Information (PI) distance and the weighted Robinson– Foulds (wRF) distance [24, 36].

### Alignment entropy calculation

To quantify the diversity and complexity of editing patterns for barcode alignments, we calculated the Shannon entropy (*H*). For a given barcode alignment, the frequency (*p*_*i*_) of each unique editing state (defined by the combination of sites) was determined relative to the total number of cells containing that barcode. The entropy is then calculated as:

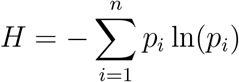

where *n* represents the total number of unique editing patterns observed for that specific barcode. Higher entropy values indicate a more diverse set of editing outcomes and higher information content, whereas lower values suggest a bias toward recurring editing states.

### Phylodynamic analysis of a monoclonal HEK293T cell culture

We applied our mechanistic model to re-analyze a dataset from a monoclonal expansion of cells from an immortalised cell line (HEK293T), originally described by [2] and previously analyzed using traditional filtering in [16].Unlike prior approaches, we utilized the original lossy barcode alignments as direct input for inference. Specifically, we randomly subsampled 1,000 cells and incorporated all 17 available barcode alignments for joint phylogenetic inference (see Fig. 14 for entropy and recovery per barcode). To account for heterogeneity in barcode recovery, the editing model was parameterized with barcode-specific heritable loss rates and dropout probabilities. The prior distributions for these parameters are detailed in Table 1.

**Table 1.** Priors used for analysis of the HEK293T dataset using the barcode loss model in SciPhy and the birth-death sampling model.

| Parameter | Distribution | 95% HDI |
| --- | --- | --- |
| Edit probabilities | Dirichlet($\alpha = 1.5$) | [1e-4, 1e-1] |
| Editing rate | Log-normal($\mu = -2, \sigma = 0.5$) | [ 0.035, 0.31 ] |
| Birth rate | Log-normal($\mu = -0.6, \sigma = 1$) | [ 0.014, 2.84 ] |
| Death rate | Log-normal($\mu = -2, \sigma = 1$) | [ 0.0035, 0.70 ] |
| Sampling proportion | 0.0008 | fixed |
| Sampling proportion | Log-normal($\mu = -7.4, \sigma = 1.2$) | [4.8e-6, 4.4e-3] |
| Origin or experiment duration | 25 | fixed |
| Heritable loss rate per barcode | Log-normal($\mu = -5, \sigma = 1$) | [0.00017, 0.034 ] |
| Dropout probability per barcode | Beta($\alpha = 3, \beta = 7$) | [ 0.054, 0.56 ] |

### Empirical analysis of mouse gastruloid development

We applied our model to DNA Typewriter barcodes from murine gastruloids [16]. This analysis consisted of two parts: (i) a re-analysis of the 780 cells and 8 barcodes from the original study to evaluate how growth estimates differ between standard filtering approaches and our explicit loss model, and (ii) an extended analysis incorporating previously discarded discarded data across three sequencing lanes (each containing two pooled biological replicates). Subsamples corresponding to individual gastruloids were first demultiplexed by hierarchical clustering of target barcode co-occurrence per lane. We then applied Shannon entropy (*H*) to identify these demultiplexed barcodes with high information content, retaining those with *H >* 2. Prior distributions for all parameters are provided in Table 2.

**Table 2.**
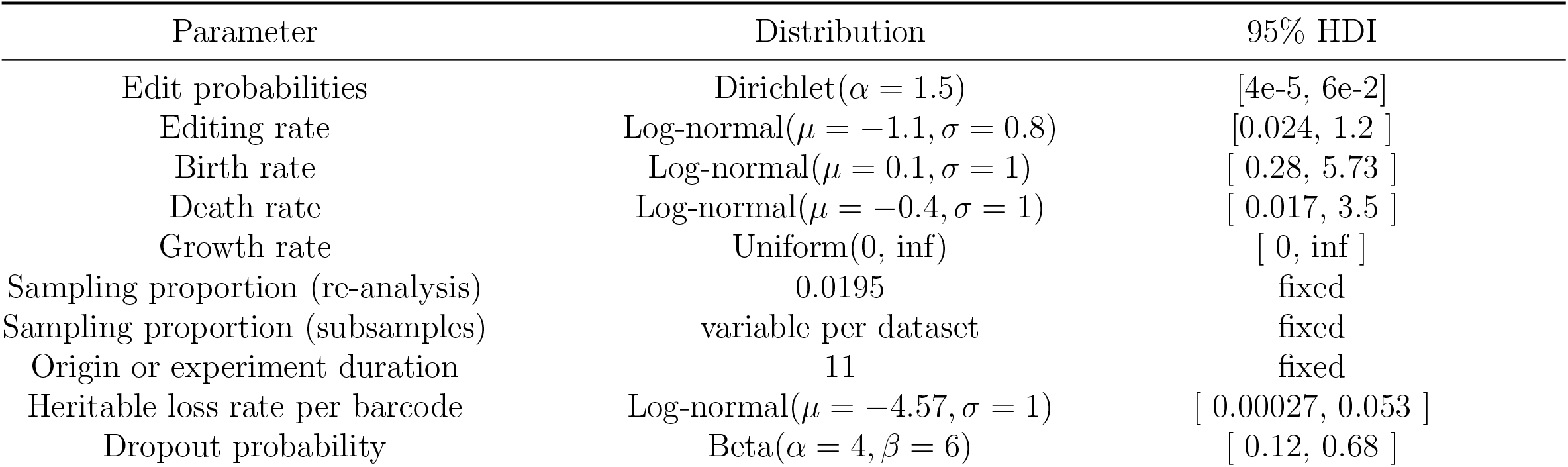
Priors used for the analysis of the gastruloid dataset using the barcode loss model in SciPhy and the birth-death sampling model.

| Parameter | Distribution | 95% HDI |
| --- | --- | --- |
| Edit probabilities | Dirichlet($\alpha = 1.5$) | [4e-5, 6e-2] |
| Editing rate | Log-normal($\mu = -1.1, \sigma = 0.8$) | [0.024, 1.2 ] |
| Birth rate | Log-normal($\mu = 0.1, \sigma = 1$) | [ 0.28, 5.73 ] |
| Death rate | Log-normal($\mu = -0.4, \sigma = 1$) | [ 0.017, 3.5 ] |
| Growth rate | Uniform(0, inf) | [ 0, inf ] |
| Sampling proportion (re-analysis) | 0.0195 | fixed |
| Sampling proportion (subsamples) | variable per dataset | fixed |
| Origin or experiment duration | 11 | fixed |
| Heritable loss rate per barcode | Log-normal($\mu = -4.57, \sigma = 1$) | [ 0.00027, 0.053 ] |
| Dropout probability | Beta($\alpha = 4, \beta = 6$) | [ 0.12, 0.68 ] |

**Table 3.**
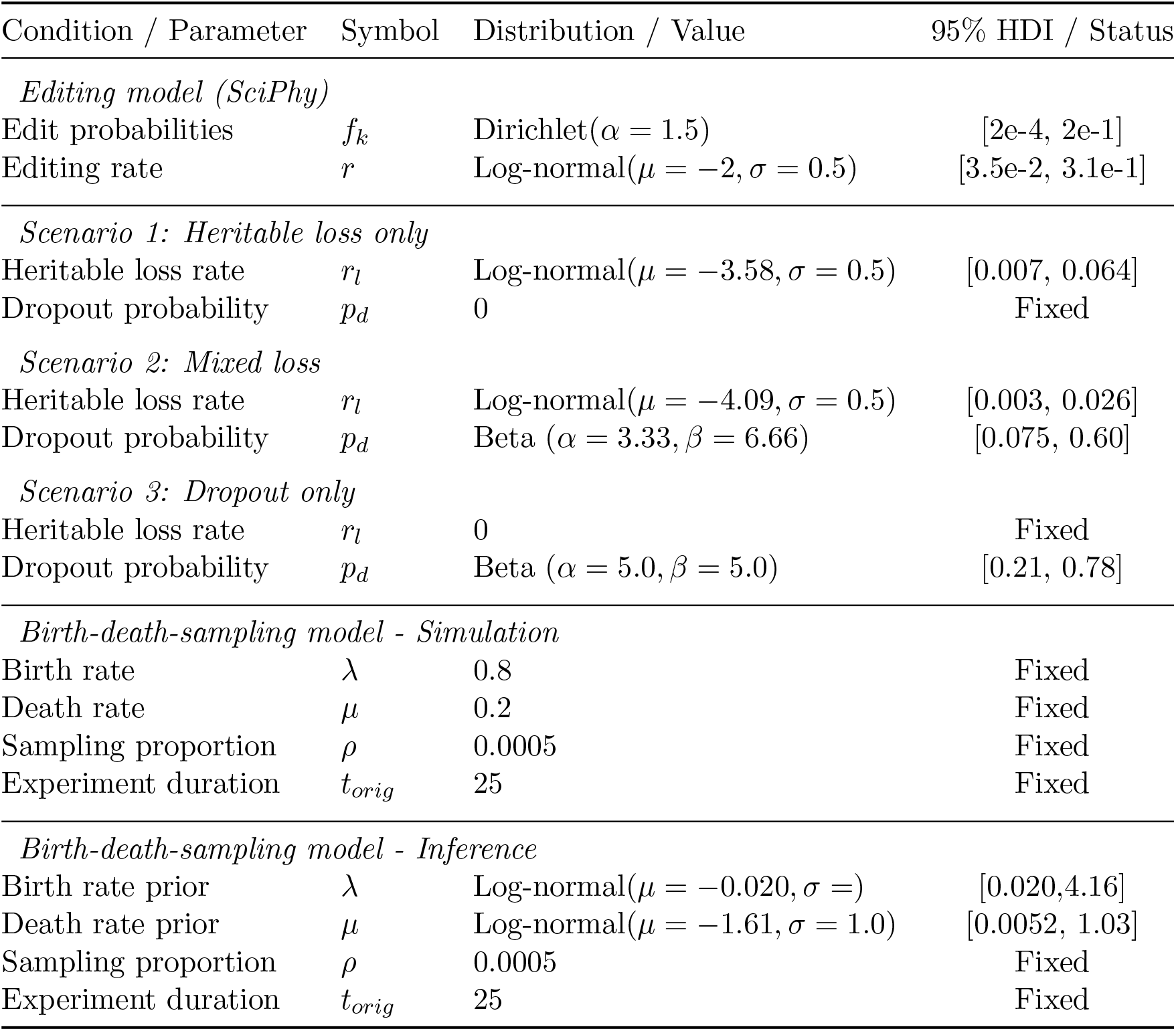
Simulation and inference parameters under low sampling (*ρ* = 0.0005) SciPhy.

**Table 4.**
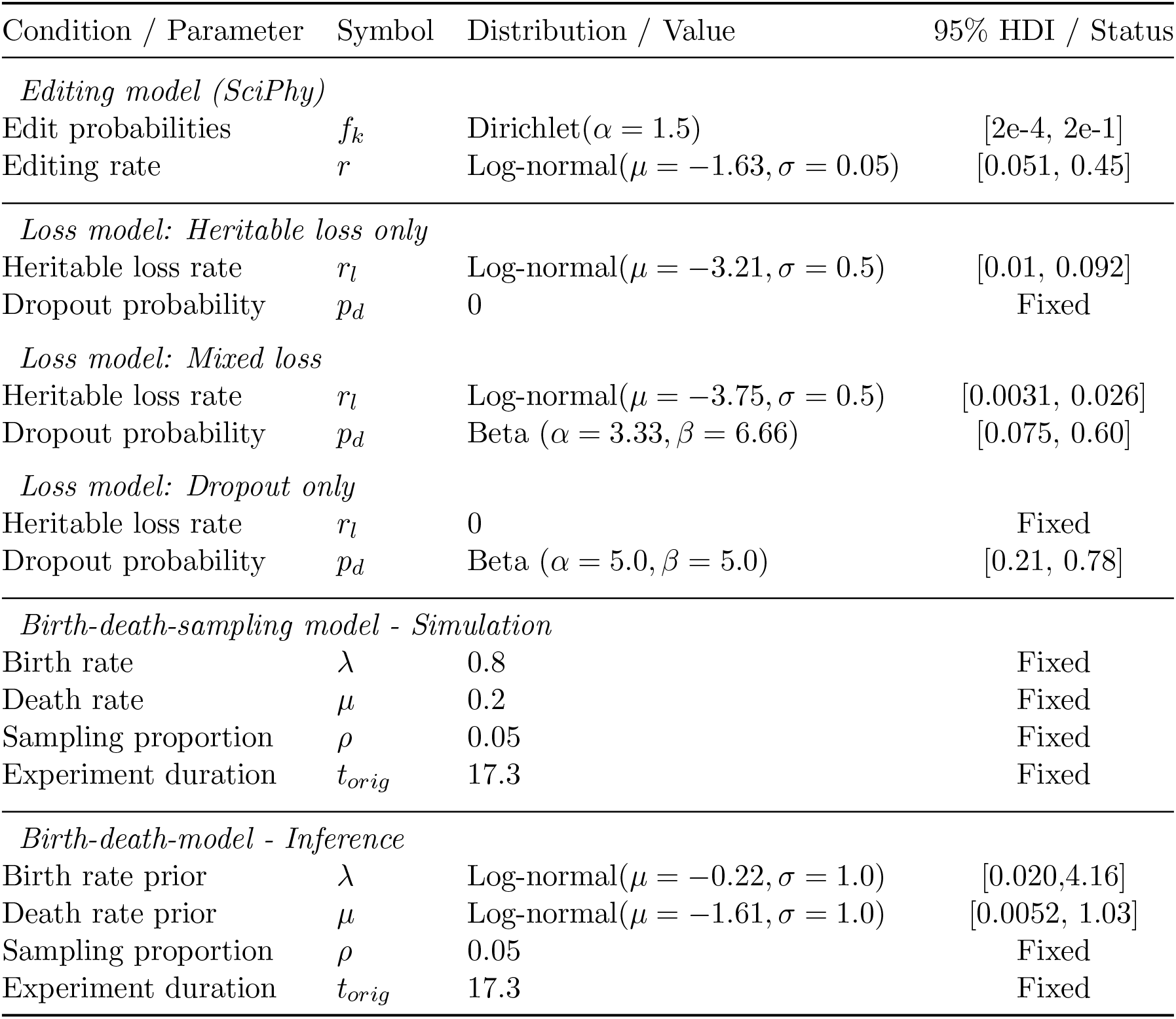
Simulation and inference parameters under intermediate sampling (*ρ* = 0.05) with the SciPhy model. The presented results for validation of the SciPhy framework extended with the loss model correspond to the “Mixed loss” scenario.

**Table 5.**
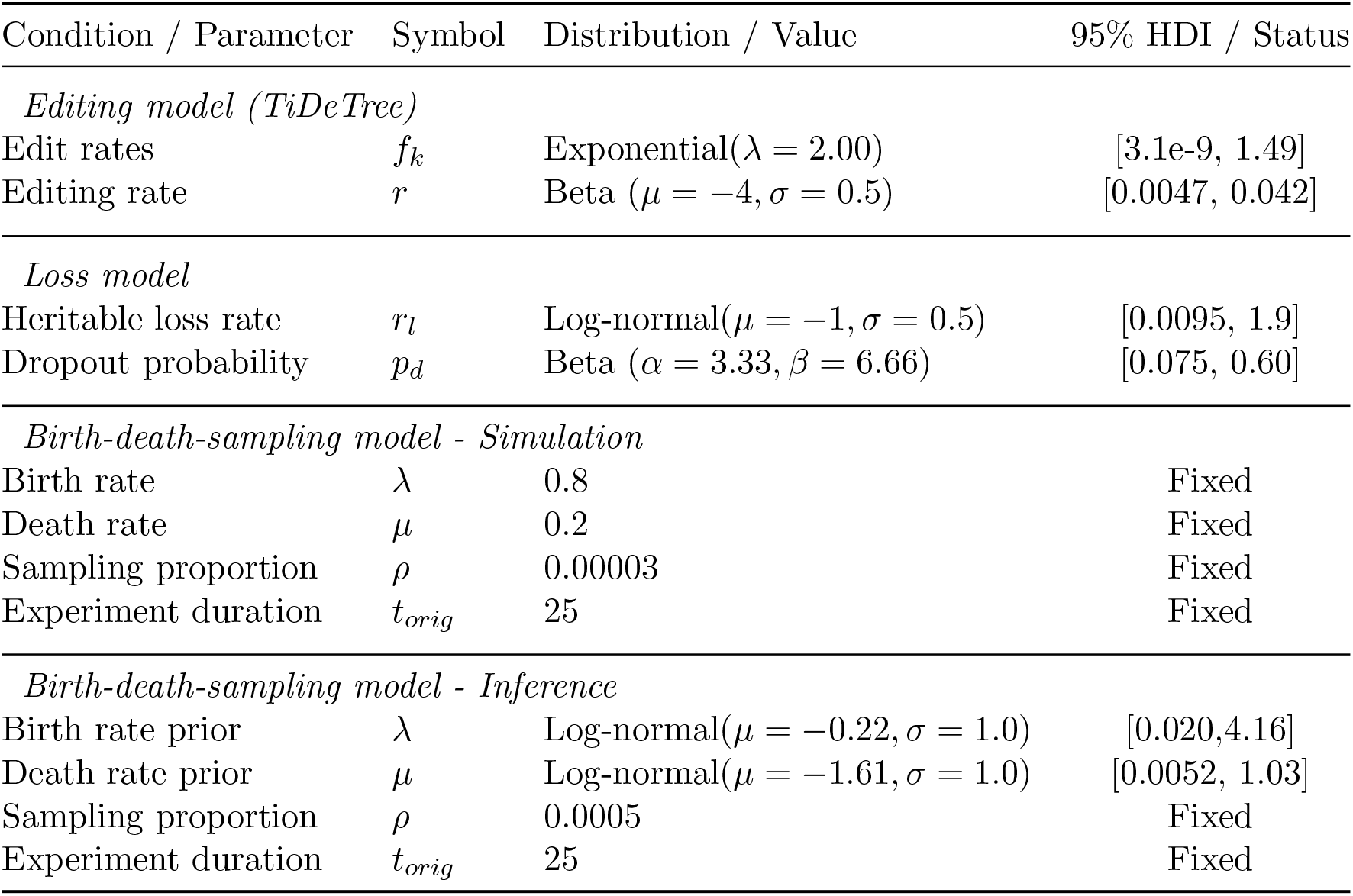
Simulation and inference priors for parameters in the validation of the TiDeTree framework extended with the loss model.

**Table 6.**
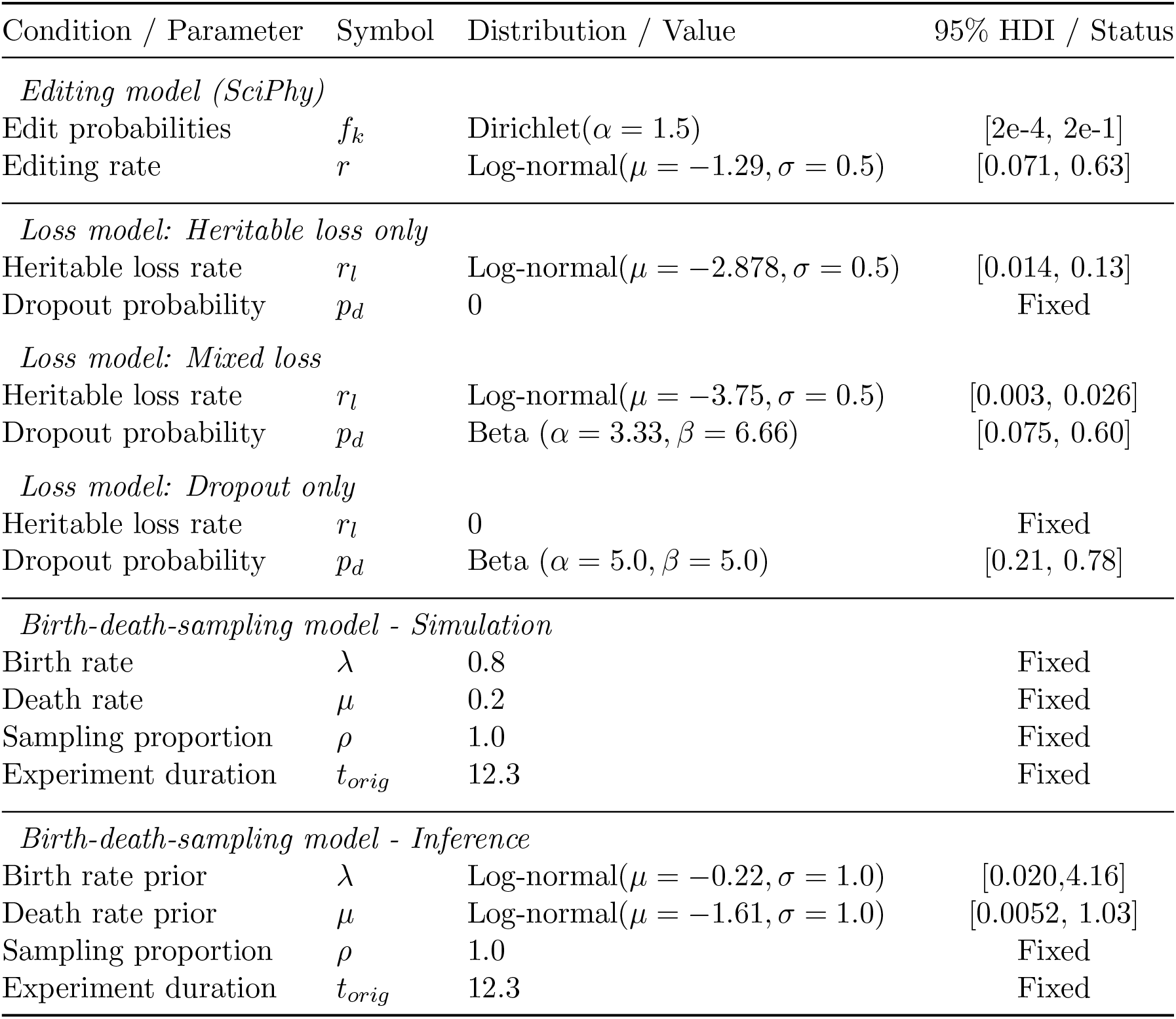
Simulation and inference parameters under high sampling (*ρ* = 1.0). SciPhy.

### Convergence of MCMC chains

For all analyses presented here, MCMC was run for as many iterations (usually in the range of 10^9^) required to reach an Effective Sample Size (ESS) of at least 200 for all parameters of interest. This criterion was used to assume the convergence of the chains. For all chains, we then discard the first 10% of the iterations, as burnin.

### Summary tree generation

For all visualizations and ground-truth comparisons, we summarize posterior tree distributions into point-estimate summary trees using the CCD0 parameterization of the conditional clade distribution (CCD) algorithm [26] in software TreeAnnotator.

## 5.1 Code availability

All code to reproduce the analyses is publicly available at https://github.com/azwaans/sciphy-error-materials. The BEAST 2 implementations are available as branches within the SciPhy and TiDeTree repositories at https://github.com/azwaans/SciPhy/tree/missing_data and https://github.com/azwaans/tidetree-dropout.

## 5.2 Data availability

Data for the monoclonal gastruloid is available under accession number GSE315827 in the Gene Expression Omnibus (GEO) database.

The data for the cell culture (HEK293T) experiment is available from following repository: https://github.com/shendurelab/DNATickerTape (see Supplementary File 2).

## 6 Competing interests

No competing interest is declared.

## 7 Author contributions statement

A.Z.: conceptualization, methodology, software, validation, formal analysis, investigation, data curation, writing—original draft, writing—review and editing, visualization and project administration. S.S.: conceptualization, methodology, writing—original draft, writing—review and editing. K.C.: conceptualization, writing—original draft, writing—review and editing. T.S.: conceptualization, methodology, resources, writing—review and editing, supervision, funding acquisition.

## 8 Acknowledgments

This project has received funding from the European Research Council (ERC) under the European Union’s Horizon 2020 research and innovation programme grant agreement no. 101001077 (to T.S.). K.C. was supported by a Marsden Fund award (22-UOA-185) from the Royal Society Te Aparāngi. S.S. was supported by a Swiss National Science Foundation (SNSF) Post-doc.Mobility fellowship (grant no. 239394). The authors thank Junhong Choi, Nicola Mulberry, Marcus Overwater, Julia Pilarski and Laura Tomás for helpful discussions.

## 9 Supplementary material

## Notes

### Competing Interest Statement

The authors have declared no competing interest.

