## Appendix for "Bayesian phylogenetics with incomplete lineage tracing data"

| Condition / Parameter | Symbol | Distribution / Value | 95% HDI / Status |
|---|---|---|---|
| *Editing model (SciPhy)* | | | |
| Edit probabilities | $f_k$ | Dirichlet($\alpha = 1.5$) | [2e-4, 2e-1] |
| Editing rate | $r$ | Log-normal($\mu = -2, \sigma = 0.5$) | [3.5e-2, 3.1e-1] |
| *Scenario 1: Heritable loss only* | | | |
| Heritable loss rate | $r_l$ | Log-normal($\mu = -3.58, \sigma = 0.5$) | [0.007, 0.064] |
| Dropout probability | $p_d$ | 0 | Fixed |
| *Scenario 2: Mixed loss* | | | |
| Heritable loss rate | $r_l$ | Log-normal($\mu = -4.09, \sigma = 0.5$) | [0.003, 0.026] |
| Dropout probability | $p_d$ | Beta ($\alpha = 3.33, \beta = 6.66$) | [0.075, 0.60] |
| *Scenario 3: Dropout only* | | | |
| Heritable loss rate | $r_l$ | 0 | Fixed |
| Dropout probability | $p_d$ | Beta ($\alpha = 5.0, \beta = 5.0$) | [0.21, 0.78] |
| *Birth-death-sampling model - Simulation* | | | |
| Birth rate | $\lambda$ | 0.8 | Fixed |
| Death rate | $\mu$ | 0.2 | Fixed |
| Sampling proportion | $\rho$ | 0.0005 | Fixed |
| Experiment duration | $t_{orig}$ | 25 | Fixed |
| *Birth-death-sampling model - Inference* | | | |
| Birth rate prior | $\lambda$ | Log-normal($\mu = -0.020, \sigma =$) | [0.020,4.16] |
| Death rate prior | $\mu$ | Log-normal($\mu = -1.61, \sigma = 1.0$) | [0.0052, 1.03] |
| Sampling proportion | $\rho$ | 0.0005 | Fixed |
| Experiment duration | $t_{orig}$ | 25 | Fixed |

**Table 1:** Simulation and inference parameters under low sampling ($\rho = 0.0005$) SciPhy.

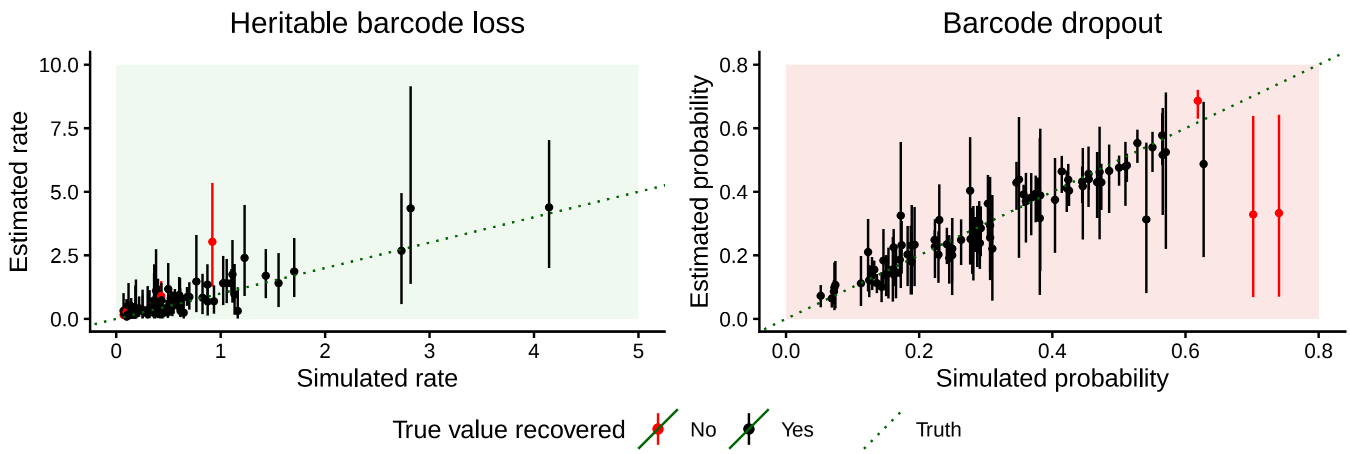


**Figure 1:** In-silico validation of the TiDeTree model extension. Validation results for theheritable loss rate (left), and dropout probability (right). Dots represent posterior medians and lines the 95% Highest Posterior Density (HPD) intervals, against a line (green) indicating the identity ($y = x$) baseline of a perfect estimator. Points and intervals are colored black if the 95% HPD interval contains the true value and red otherwise. The model achieves coverage of 93%, 96%, and 94% for the editing rate, heritable loss rate, and dropout probability, respectively.

| Condition / Parameter | Symbol | Distribution / Value | 95% HDI / Status |
|---|---|---|---|
| *Editing model (SciPhy)* | | | |
| Edit probabilities | $f_k$ | Dirichlet($\alpha = 1.5$) | [2e-4, 2e-1] |
| Editing rate | $r$ | Log-normal($\mu = -1.63, \sigma = 0.05$) | [0.051, 0.45] |
| *Loss model: Heritable loss only* | | | |
| Heritable loss rate | $r_l$ | Log-normal($\mu = -3.21, \sigma = 0.5$) | [0.01, 0.092] |
| Dropout probability | $p_d$ | 0 | Fixed |
| *Loss model: Mixed loss* | | | |
| Heritable loss rate | $r_l$ | Log-normal($\mu = -3.75, \sigma = 0.5$) | [0.0031, 0.026] |
| Dropout probability | $p_d$ | Beta ($\alpha = 3.33, \beta = 6.66$) | [0.075, 0.60] |
| *Loss model: Dropout only* | | | |
| Heritable loss rate | $r_l$ | 0 | Fixed |
| Dropout probability | $p_d$ | Beta ($\alpha = 5.0, \beta = 5.0$) | [0.21, 0.78] |
| *Birth-death-sampling model - Simulation* | | | |
| Birth rate | $\lambda$ | 0.8 | Fixed |
| Death rate | $\mu$ | 0.2 | Fixed |
| Sampling proportion | $\rho$ | 0.05 | Fixed |
| Experiment duration | $t_{orig}$ | 17.3 | Fixed |
| *Birth-death-model - Inference* | | | |
| Birth rate prior | $\lambda$ | Log-normal($\mu = -0.22, \sigma = 1.0$) | [0.020,4.16] |
| Death rate prior | $\mu$ | Log-normal($\mu = -1.61, \sigma = 1.0$) | [0.0052, 1.03] |
| Sampling proportion | $\rho$ | 0.05 | Fixed |
| Experiment duration | $t_{orig}$ | 17.3 | Fixed |

**Table 2:** Simulation and inference parameters under intermediate sampling ($\rho = 0.05$) with the SciPhy model. The presented results for validation of the SciPhy framework extended with the loss model correspond to the "Mixed loss" scenario.

| Condition / Parameter | Symbol | Distribution / Value | 95% HDI / Status |
|---|---|---|---|
| *Editing model (TiDeTree)* | | | |
| Edit rates | $f_k$ | Exponential($\lambda = 2.00$) | [3.1e-9, 1.49] |
| Editing rate | $r$ | Beta ($\mu = -4, \sigma = 0.5$) | [0.0047, 0.042] |
| *Loss model* | | | |
| Heritable loss rate | $r_l$ | Log-normal($\mu = -1, \sigma = 0.5$) | [0.0095, 1.9] |
| Dropout probability | $p_d$ | Beta ($\alpha = 3.33, \beta = 6.66$) | [0.075, 0.60] |
| *Birth-death-sampling model - Simulation* | | | |
| Birth rate | $\lambda$ | 0.8 | Fixed |
| Death rate | $\mu$ | 0.2 | Fixed |
| Sampling proportion | $\rho$ | 0.00003 | Fixed |
| Experiment duration | $t_{orig}$ | 25 | Fixed |
| *Birth-death-sampling model - Inference* | | | |
| Birth rate prior | $\lambda$ | Log-normal($\mu = -0.22, \sigma = 1.0$) | [0.020,4.16] |
| Death rate prior | $\mu$ | Log-normal($\mu = -1.61, \sigma = 1.0$) | [0.0052, 1.03] |
| Sampling proportion | $\rho$ | 0.0005 | Fixed |
| Experiment duration | $t_{orig}$ | 25 | Fixed |

**Table 3:** Simulation and inference priors for parameters in the validation of the TiDeTree framework extended with the loss model.

| Condition / Parameter | Symbol | Distribution / Value | 95% HDI / Status |
|---|---|---|---|
| *Editing model (SciPhy)* | | | |
| Edit probabilities | $f_k$ | Dirichlet($\alpha = 1.5$) | [2e-4, 2e-1] |
| Editing rate | $r$ | Log-normal($\mu = -1.29, \sigma = 0.5$) | [0.071, 0.63] |
| *Loss model: Heritable loss only* | | | |
| Heritable loss rate | $r_l$ | Log-normal($\mu = -2.878, \sigma = 0.5$) | [0.014, 0.13] |
| Dropout probability | $p_d$ | 0 | Fixed |
| *Loss model: Mixed loss* | | | |
| Heritable loss rate | $r_l$ | Log-normal($\mu = -3.75, \sigma = 0.5$) | [0.003, 0.026] |
| Dropout probability | $p_d$ | Beta ($\alpha = 3.33, \beta = 6.66$) | [0.075, 0.60] |
| *Loss model: Dropout only* | | | |
| Heritable loss rate | $r_l$ | 0 | Fixed |
| Dropout probability | $p_d$ | Beta ($\alpha = 5.0, \beta = 5.0$) | [0.21, 0.78] |
| *Birth-death-sampling model - Simulation* | | | |
| Birth rate | $\lambda$ | 0.8 | Fixed |
| Death rate | $\mu$ | 0.2 | Fixed |
| Sampling proportion | $\rho$ | 1.0 | Fixed |
| Experiment duration | $t_{orig}$ | 12.3 | Fixed |
| *Birth-death-sampling model - Inference* | | | |
| Birth rate prior | $\lambda$ | Log-normal($\mu = -0.22, \sigma = 1.0$) | [0.020,4.16] |
| Death rate prior | $\mu$ | Log-normal($\mu = -1.61, \sigma = 1.0$) | [0.0052, 1.03] |
| Sampling proportion | $\rho$ | 1.0 | Fixed |
| Experiment duration | $t_{orig}$ | 12.3 | Fixed |

**Table 4:** Simulation and inference parameters under high sampling ($\rho = 1.0$). SciPhy

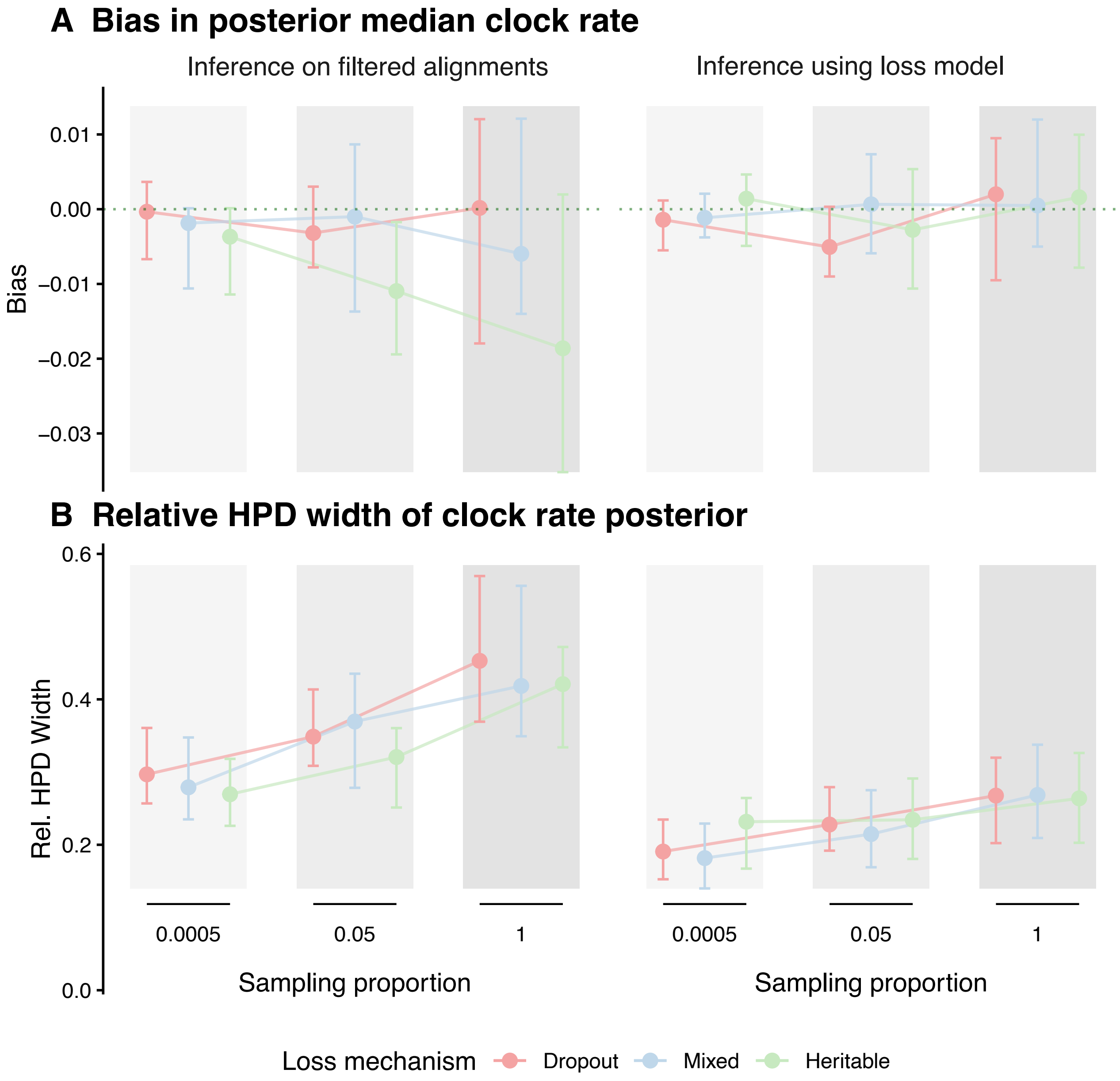


**Figure 2:** Simulation-based comparison of editing (clock) rate estimation under data filtering and mechanistic loss modeling. **A** Absolute bias and **B** Mean relative HPD width The estimation is compared varying loss scenarios (colors) and tree shapes (modulated by sampling proportion, background shading). Points represent the mean performance across replicates, with error bars denoting the 95% interval of the estimates. In A, the horizontal dotted line indicates an ideal bias of zero.

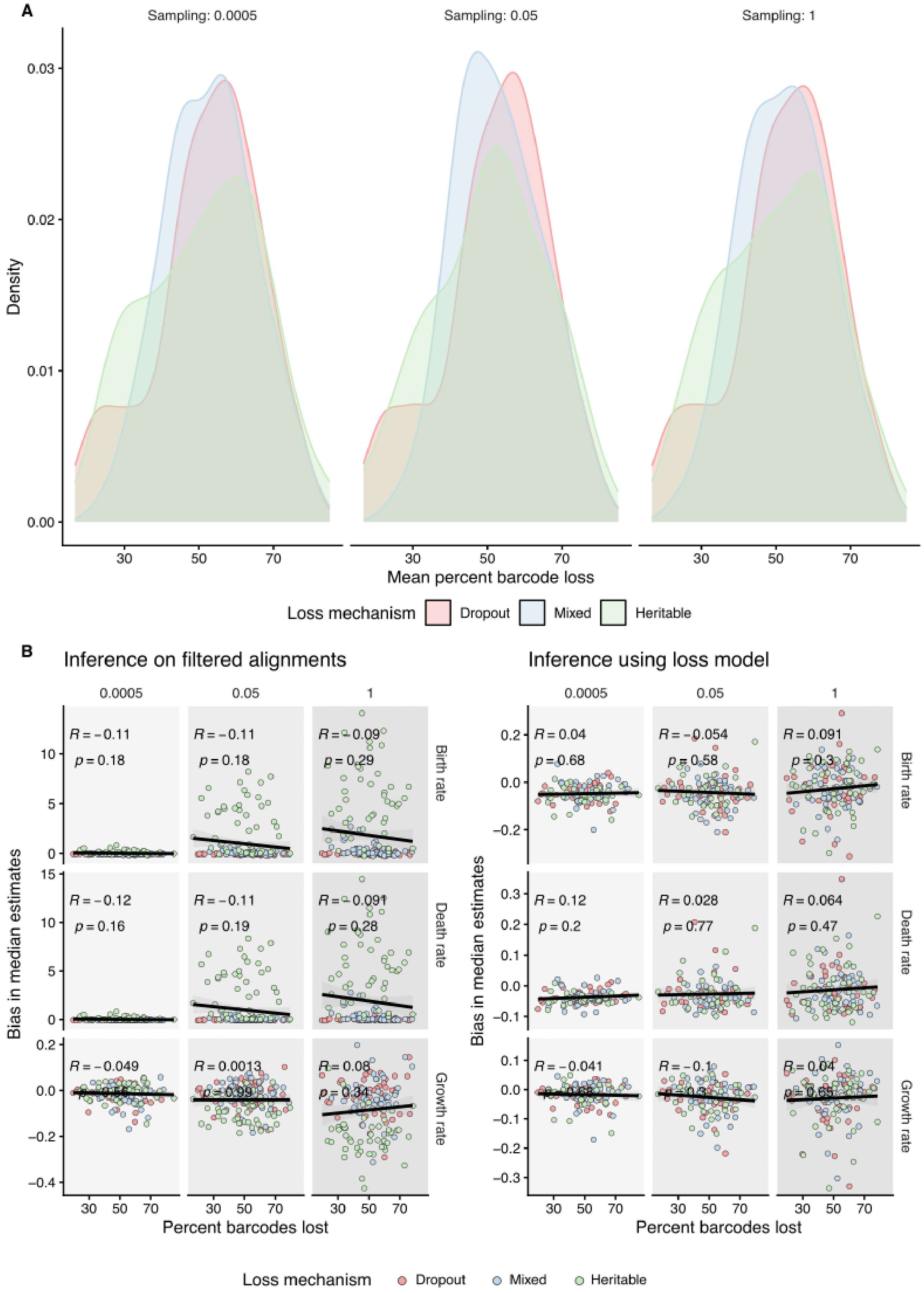


**Figure 3:** Impact of overall barcode loss on parameter bias. **A** Density distributions of the mean percentage of barcode loss over all simulated datasets across sampling proportions (columns) and loss mechanisms (colors). Total data loss remains consistent at approximately 50% across all scenarios, isolating the effect of the loss mechanism rather than the loss magnitude. **B** Correlation between the percentage of data loss and bias in birth ($\lambda$), death ($\mu$), and net growth ($r$) rates.

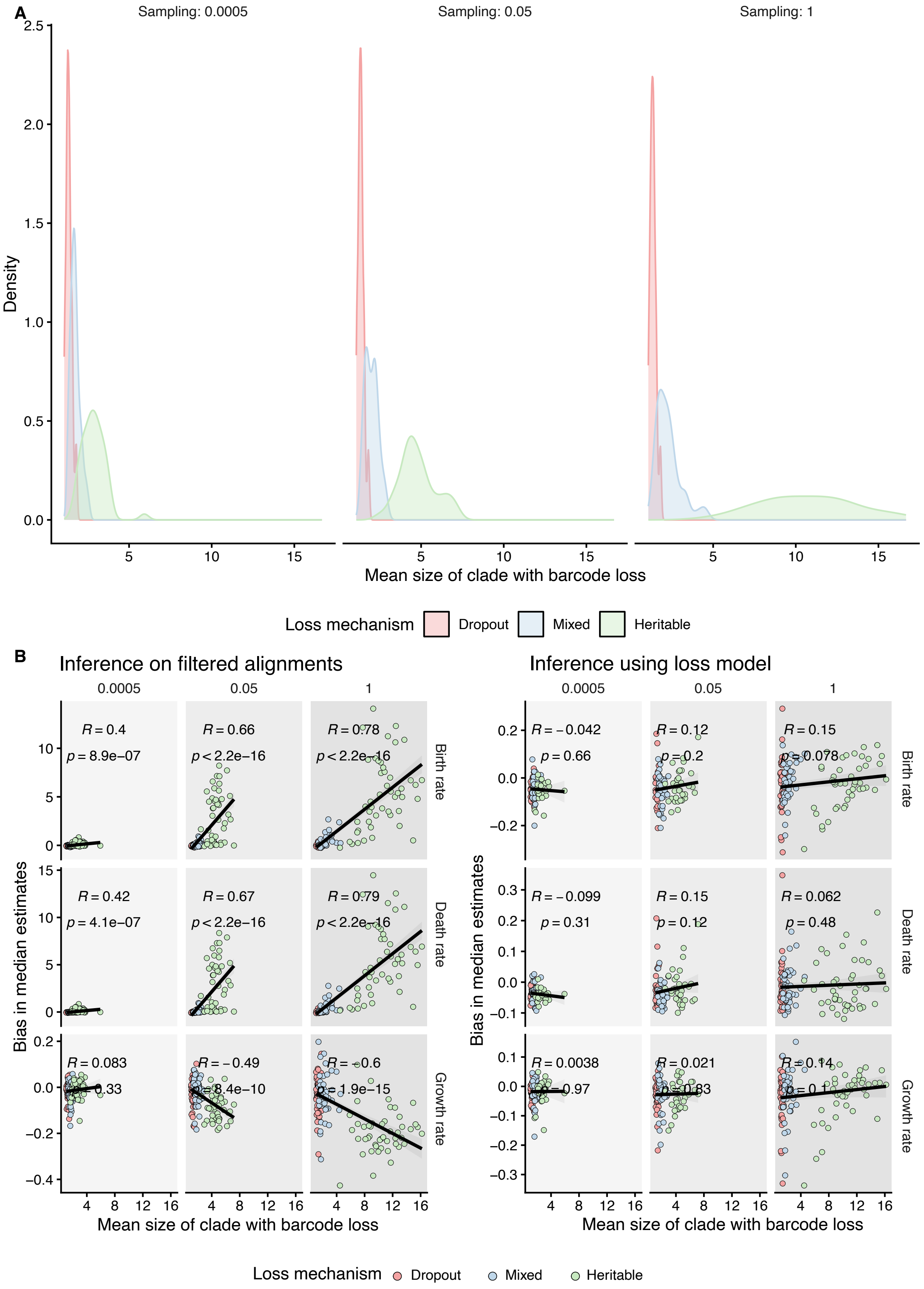


**Figure 4:** Impact of lost clade size on parameter bias. **A** Density distributions showing the mean size of clades affected by data loss over all simulated datasets across the three sampling proportions (columns) and three loss mechanisms (colors). Heritable loss (green) results in larger monophyletic clades with lost barcodes compared to dropout (pink), a distinction that intensifies as sampling density increases. B Correlation between lost clade size and bias in birth, death, and net growth, rates.

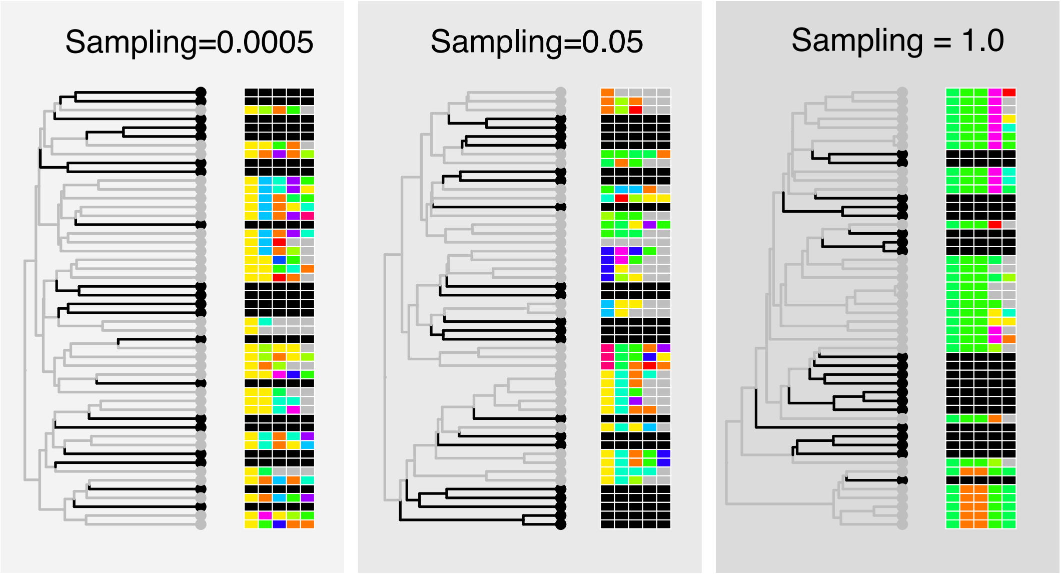


**Figure 5:** Explanatory schematic illustrating the effect of sampling proportion on tree branching and heritable barcode loss patterns. Holding total tree size and heritable loss rate constant, increasing the sampling proportion (left to right) shifts branching toward the root. This topological shift causes heritable loss events to affect larger, highly clustered clades, resulting in increasingly aggregated missing data across the alignment.

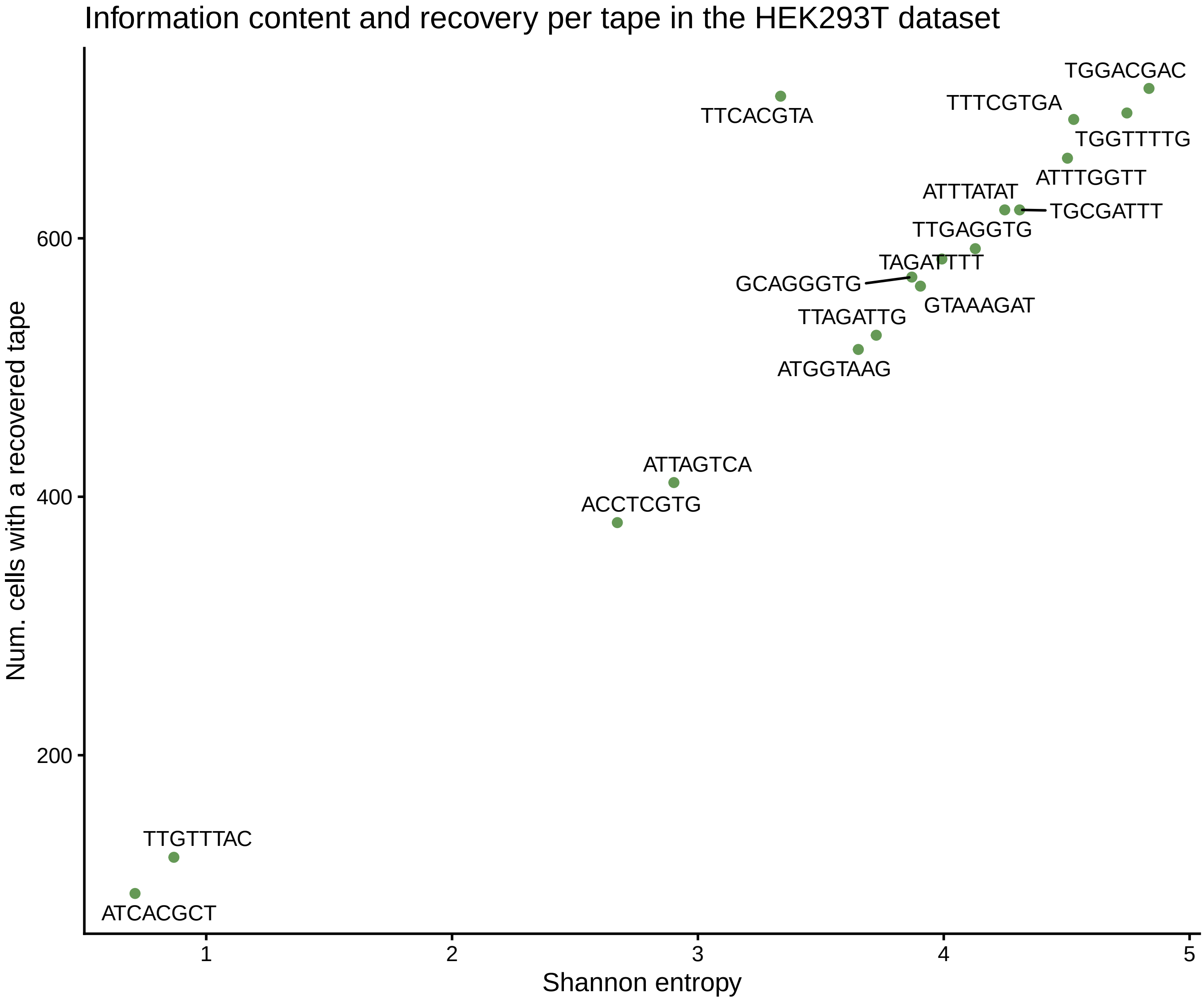


**Figure 6:** Information content and barcode recovery per tape in the HEK293T dataset. Scatter plot showing the relationship between Shannon entropy (x-axis) and cell recovery (y-axis) for each target barcode or "tape" used in the dataset.

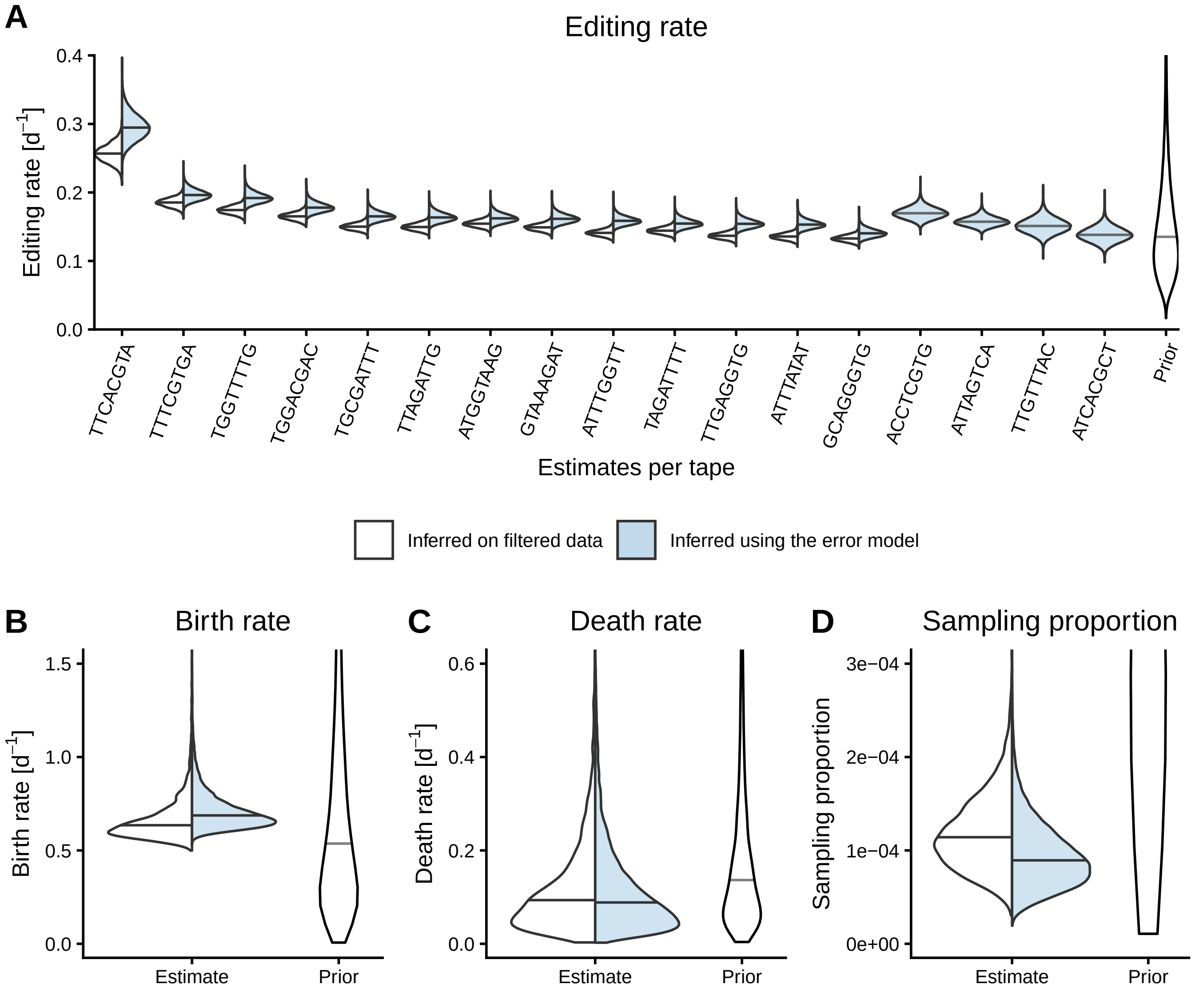


**Figure 7:** Comparison of phylogenetic estimates obtained in SciPhy for the monoclonal growth of HEK293T cells harboring DNA Typewriter barcodes, when filtering out lossy cells or with our model extension co-estimating the sampling proportion. We show estimates of the editing rate per barcode (**A**) and birth and death rates (**B** and **C**) estimated using both approaches.

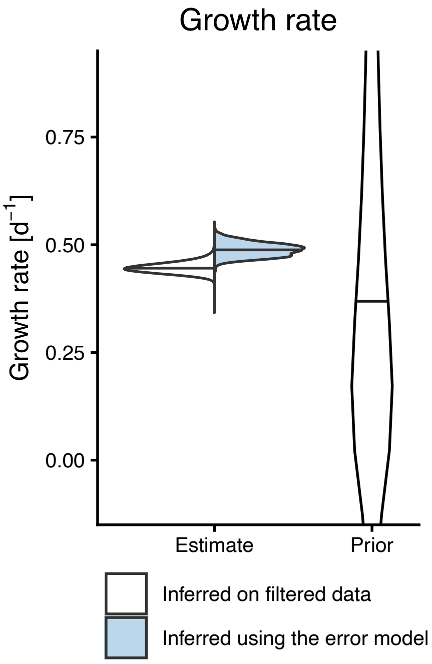


**Figure 8:** Comparison of estimates of the growth rate obtained in SciPhy for the monoclonal growth of HEK293T cells harboring DNA Typewriter barcodes, when filtering out lossy cells or with our model extension, with the sampling proportion fixed to 0.0008.

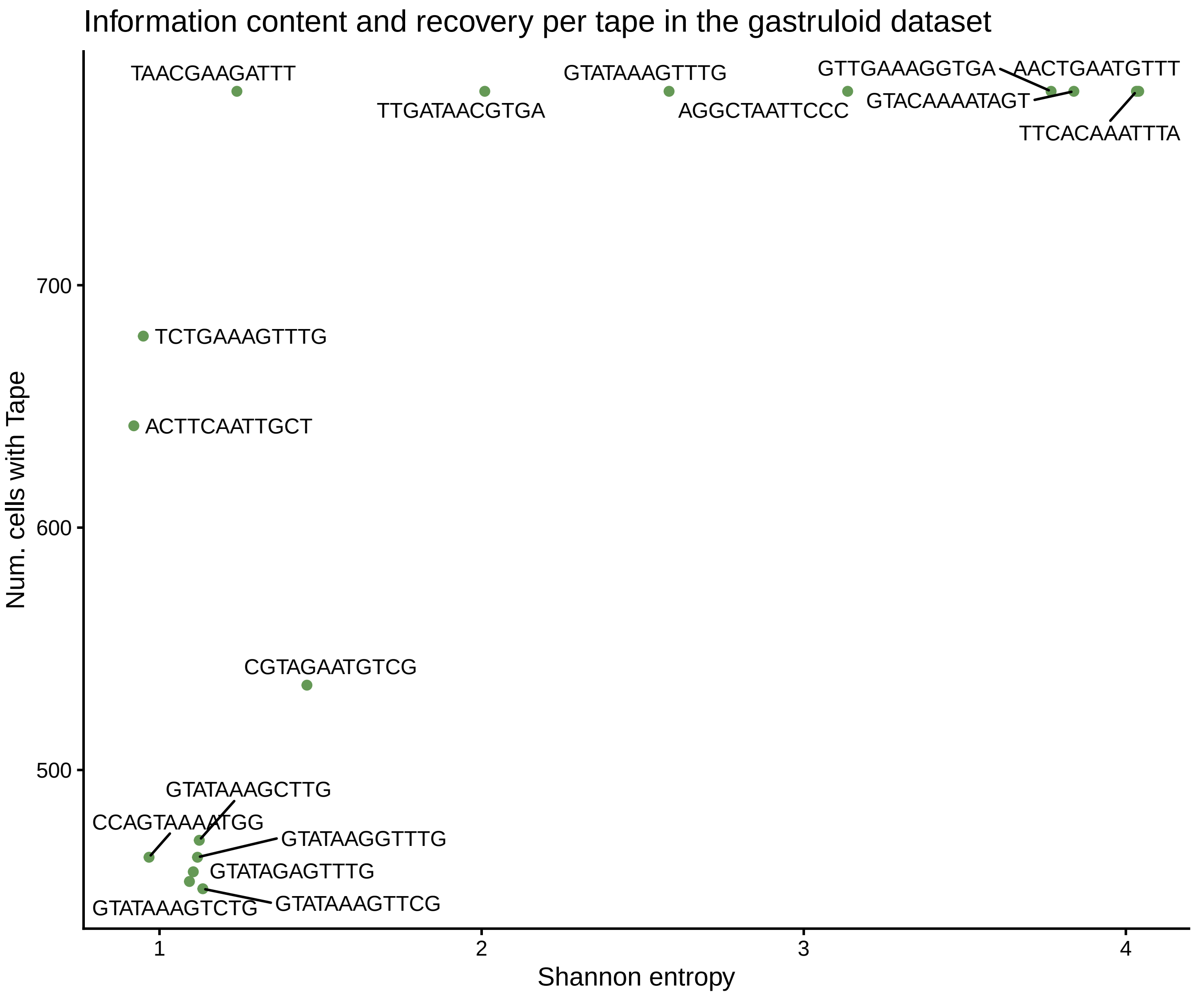


**Figure 9:** Information content and barcode recovery per tape in the gastruloid dataset analysed in [1]. Scatter plot showing the relationship between Shannon entropy (x-axis) and cell recovery (y-axis) for each target barcode or "tape" used in the dataset.

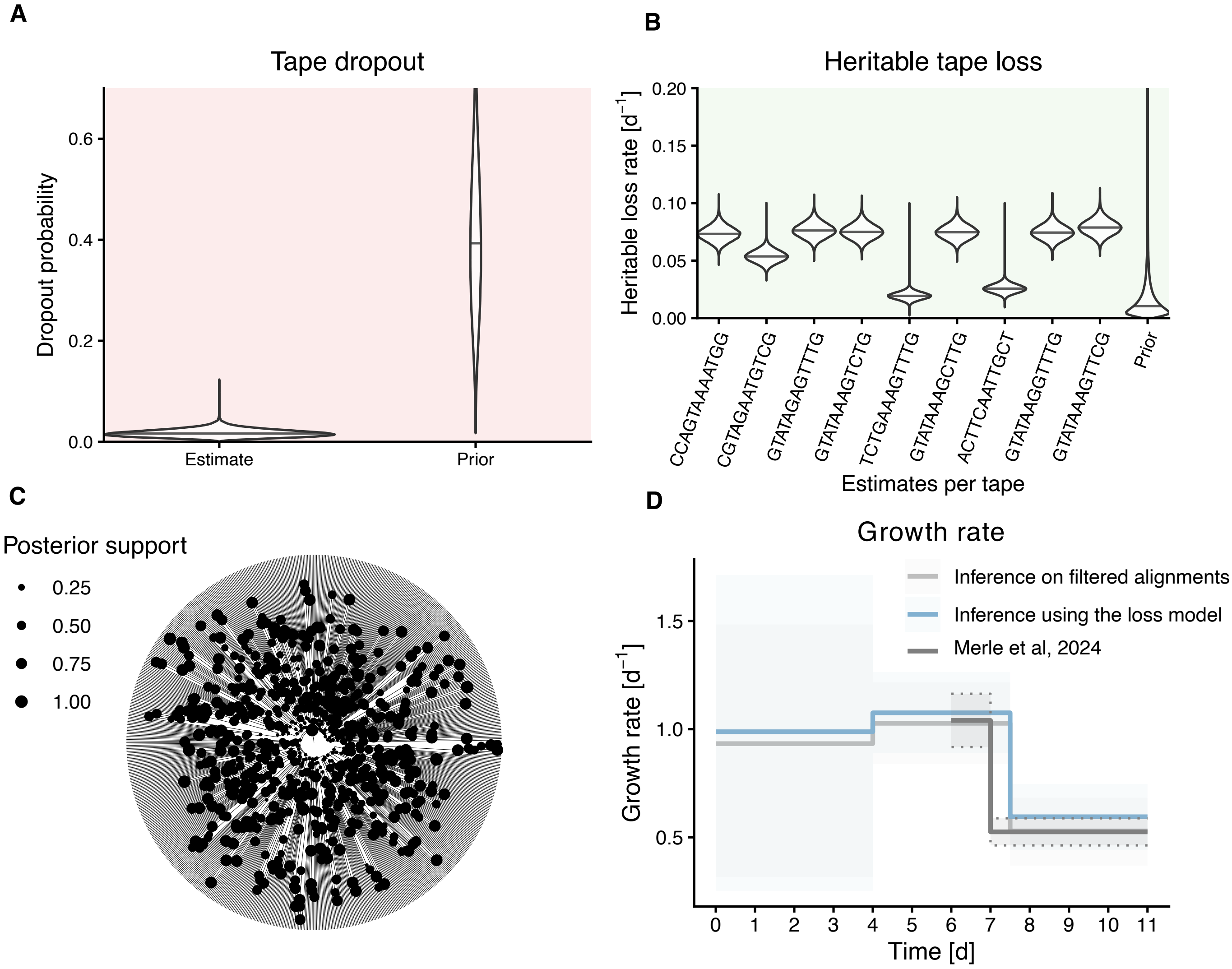


**Figure 10:** Comparison of phylogenetic estimates obtained in SciPhy for the growth of mouse gastruloid from a founding mouse embryonic stem cell (mESC) cell harboring DNA Typewriter barcodes, when filtering out lossy cells or with our model extension. In **A** and **B** we show estimates of the editing rate per barcode and growth rate estimated using both approaches. In **C** and **D** we show estimates of the heritable loss rate per barcode, and dropout probability using the loss model.

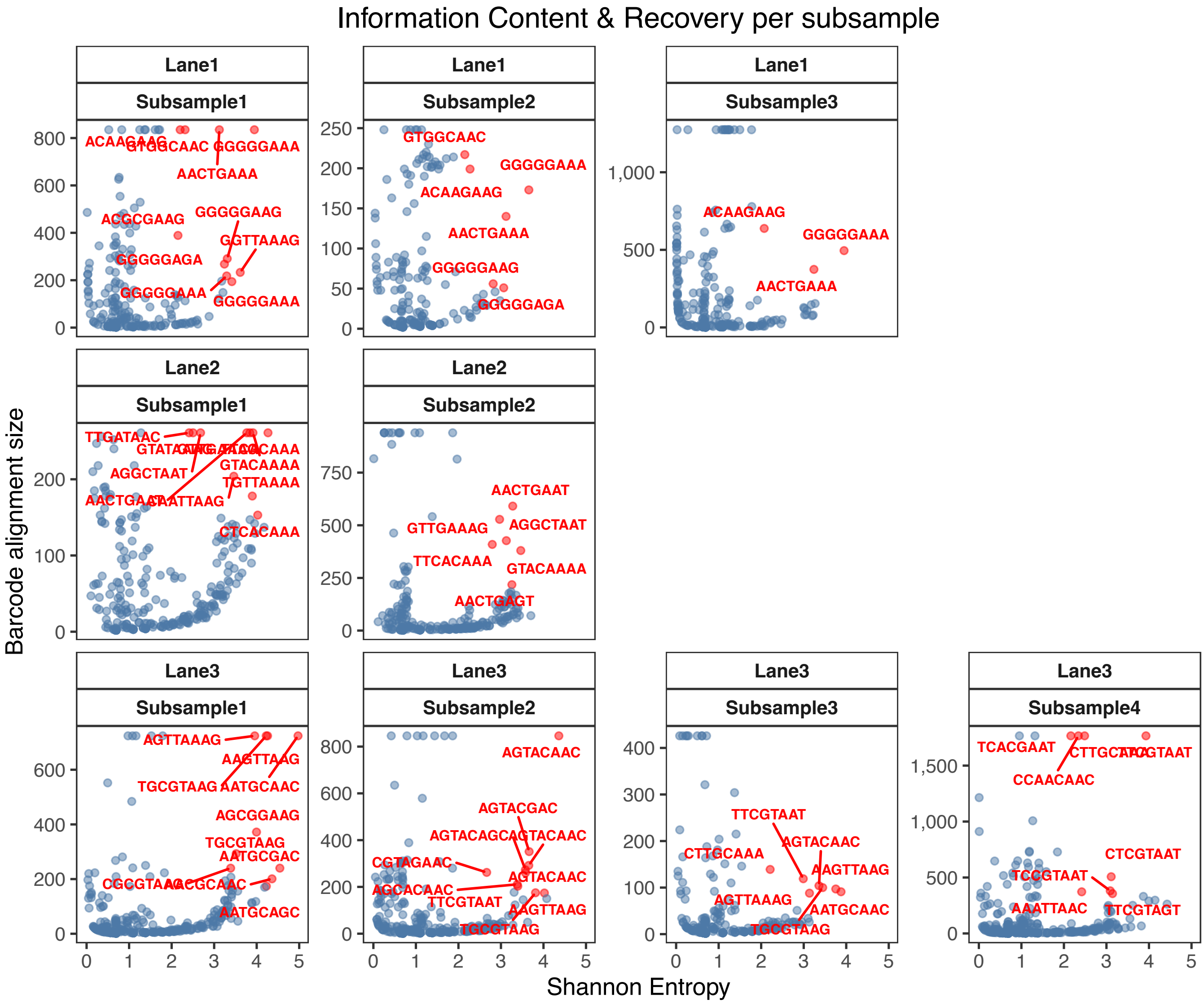


**Figure 11:** Information content and barcode recovery across subsamples of gastruloid replicates. Scatter plot shows the relationship between Shannon entropy ($x$-axis) and total cell recovery ($y$-axis) for every unique target barcode ("tape") across all subsamples. Red highlights indicate specific barcodes included for analysis (with Shannon entropy ¿ 2 and recovery in more than 20% of cells) with labels corresponding to their identifiers. Facets are organized by experimental lane and subsample. The subsamples included in main are the top 3 datasets by mean alignment entropy, Lane 2 and 1 subsamples 1 and 2 and Lane 3 subsample 1

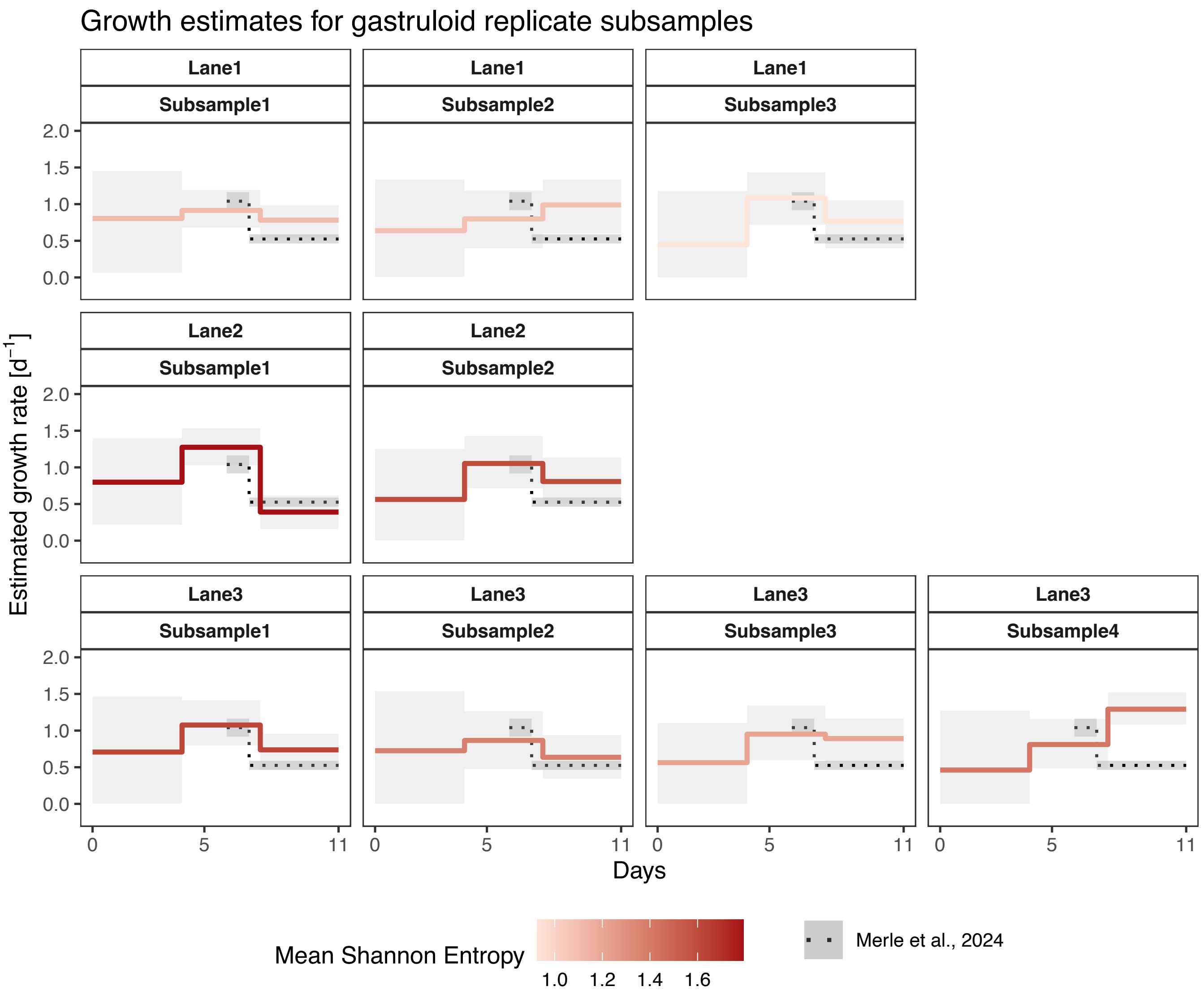


**Figure 12:** Reconstructed growth dynamics across subsamples of gastruloid replicates. Inferred growth rates obtained using the SciPhy framework, incorporating a barcode loss model under a birth-death-sampling process. Data represent subsampled populations derived from replicates of gastruloids derived from mouse embryonic stem cells (mESCs) harboring DNA Typewriter barcodes. Each panel displays the median posterior growth rate (solid colored lines) and 95% HPD intervals (grey ribbons), with lines colored according to the mean Shannon entropy of the corresponding dataset. The black dotted line and shaded region represent reference growth data from Merle et al. (2024). The subsamples included in main are the top 3 datasets by mean alignment entropy, Lane 2 and 1 subsamples 1 and 2 and Lane 3 subsample 1.

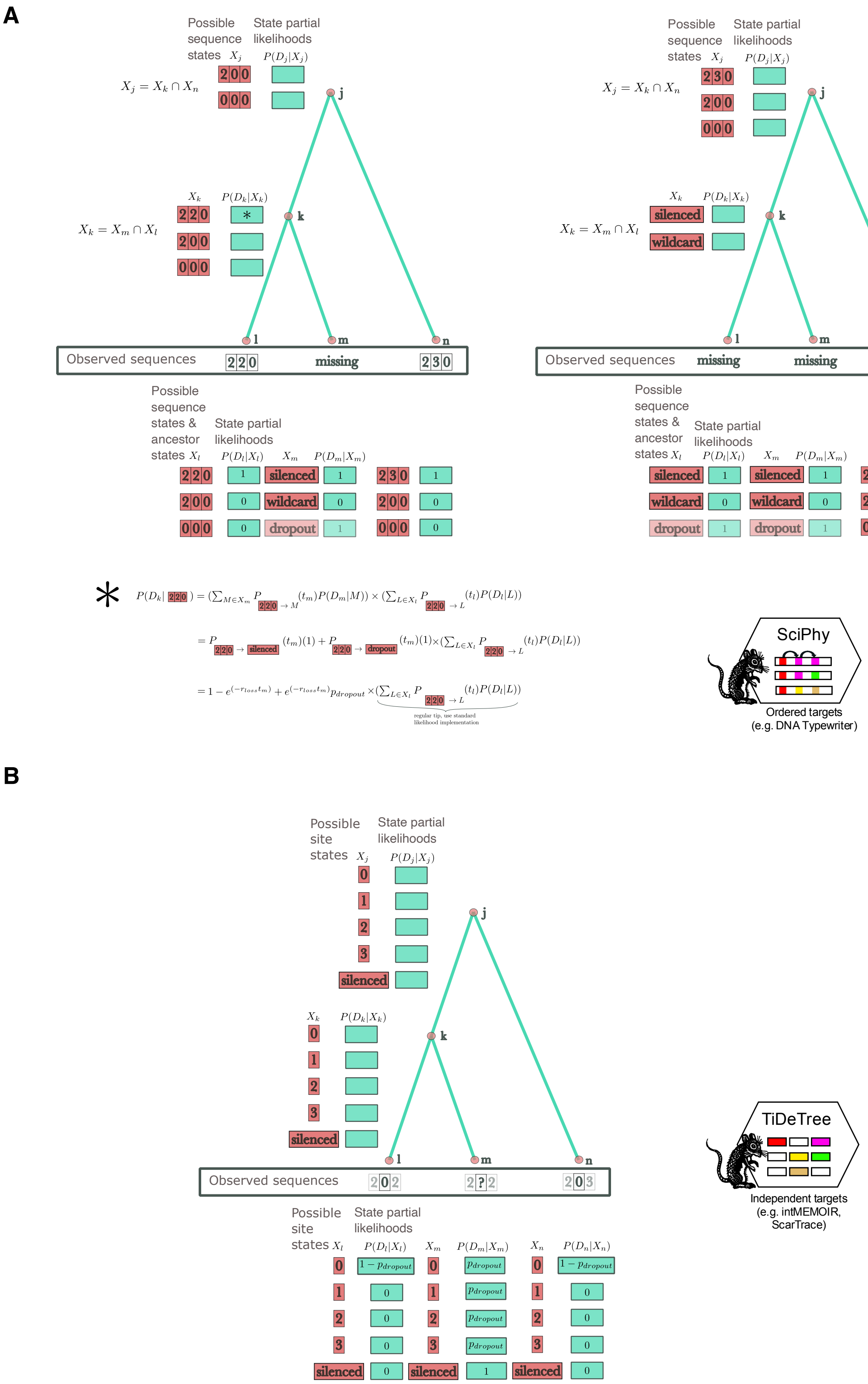


**Figure 13:** Integrating the loss model into the likelihood implementations of SciPhy and TiDeTree. **A** SciPhy: Partial likelihoods are recursively calculated for sets of barcode sequence states, determined at each node as the intersection of descendant state sets (excluding the dropout state, which can only appear at tips). We extended this framework to accommodate missing data by introducing a dropout state, a silenced state, and a wildcard state (representing any valid barcode sequence state that was neither silenced nor dropped out). Two examples illustrate how missing barcode sequences dictate ancestral states at internal nodes, alongside an example of the resulting partial likelihood calculation. **B** TiDeTree: Conversely, TiDeTree calculates partial likelihoods for individual sites within the barcodes, allowing it to operate on a fixed state space of all possible characters at each node—which already included a silenced state. Consequently, the TiDeTree implementation is extended by directly factoring the dropout probability into the site-specific partial likelihoods at the tips, as shown in the example.
